# Human heart assembloids with autologous tissue-resident macrophages recreate physiological immuno-cardiac interactions

**DOI:** 10.1101/2024.11.13.623051

**Authors:** Colin O’Hern, Sammantha Caywood, Shakhlo Aminova, Artem Kiselev, Brett Volmert, Fei Wang, Merlinda-Loriane Sewavi, Weiheng Cao, Mia Dionise, Priyadharshni Muniyandi, Mirel Popa, Hussain Basrai, Milana Skoric, George Boulos, Amanda Huang, Isabel Nuñez-Regueiro, Nagib Chalfoun, Sangbum Park, Nureddin Ashammakhi, Chao Zhou, Christopher Contag, Aitor Aguirre

**Affiliations:** Institute for Quantitative Health Science and Engineering, Michigan State University, East Lansing, MI, USA; Department of Biomedical Engineering, Michigan State University, East Lansing, MI, USA; College of Osteopathic Medicine, Michigan State University, East Lansing, MI, USA; Department of Medicine, Michigan State University, East Lansing, MI, USA; Department of Pharmacology and Toxicology, Michigan State University, East Lansing, MI, USA; Department of Biomedical Engineering, McKelvey School of Engineering, Washington University in St. Louis, St. Louis, MO, USA; Institute for Cellular Biology and Pathology “Nicolae Simionescu” of the Romanian Academy, Bucharest, Romania; Frederick Meijer Heart and Vascular Institute, Corewell Health, Grand Rapids, MI, USA; Department of Microbiology, Genetics and Immunology, Michigan State University, East Lansing, MI, USA

**Keywords:** human heart organoid, assembloid, pluripotent stem cell, atrial fibrillation, cardiac development, inflammasome

## Abstract

Interactions between the developing heart and the embryonic immune system are essential for proper cardiac development and maintaining homeostasis, with disruptions linked to various diseases. While human pluripotent stem cell (hPSC)-derived organoids are valuable models for studying human organ function, they often lack critical tissue-resident immune cells. Here, we introduce an advanced human heart assembloid model, termed hHMA (human heart-macrophage assembloid), which fully integrates autologous cardiac tissue-resident macrophages (MPs) with pre-existing human heart organoids (hHOs). Through multi-omic analyses, we confirmed that these MPs are phenotypically similar to embryonic cardiac tissue-resident MPs and remain viable in the assembloids over time. The inclusion of MPs significantly impacts hHMA development, influencing cardiac cellular composition, boosting cellular communication, remodeling the extracellular matrix, promoting ventricular morphogenesis, and enhancing sarcomeric maturation. Our findings indicate that MPs contribute to homeostasis via efferocytosis, integrate into the cardiomyocyte electrical system, and support catabolic metabolism. To demonstrate the versatility of this model, we developed a platform to study cardiac arrhythmias by chronic exposure to pro-inflammatory factors linked to arrhythmogenesis in clinical settings, successfully replicating key features of inflammasome-mediated atrial fibrillation. Overall, this work introduces a robust platform for examining the role of immune cells in cardiac development, disease mechanisms, and drug discovery, bridging the gap between *in vitro* models and human physiology. These findings offer insights into cardiogenesis and inflammation-driven heart disease, positioning the hHMA system as an invaluable tool for future cardiovascular research and therapeutic development.

## INTRODUCTION

Cardiovascular diseases are the leading cause of death in developed countries^1,2^. Despite a need to study human cardiovascular development and pathology, most of our knowledge about the of the human heart is based on animal models^3,4^ that do not recapitulate many aspects of the human heart^5–7^. This is particularly limiting when studying human heart development, where direct access to human tissue is restricted, but it also applies to adult conditions in many instances. To overcome these research limitations, significant efforts have been made to engineer human heart models *in vitro* that are faithful to the heart’s physiology^8–10^. With the advent of hPSCs^11,12^ protocols have been developed to generate more sophisticated cardiac cell types in culture, for disease modeling, drug studies, and other applications^13–15^. However, these models frequently fall short in recapitulating the complexity of the human heart. To create more complex heart tissues, approaches based on cellular self-assembly, bioengineering techniques, or a combination of both^16^ have been developed to create different types of cardiac organoids. Among cardiac organoid models, those that rely on cellular self-organization have recently shown great promise^17–21^, due to their ability to mimic the heart’s structural organization and multicellular composition. Our group^17,22,23^ has developed advanced hHO systems derived from hPSCs through different strategies. These hHOs recapitulate human heart development, including anatomical features (chambers, layering, proepicardial organ), function, cellular complexity (ventricular and atrial cardiomyocytes, cardiac fibroblasts, endocardial cells, endothelial cells, conductance cells, epicardial cells), and development, including heart tube patterning. Although the hHOs mimic many aspects of the developing human heart, they still lack some key cell types in heart development that migrate from other sites into the developing heart, such as tissue-resident MPs.

Most of what we know about tissue-resident MPs during heart development is also based on animal models^24–25^. In mice, yolk sac-derived MPs first appear shortly after the heart begins to beat^24^. A few days later, MPs derived from the fetal liver seed the heart^26^, and once bone calcification begins, the remainder of MPs are recruited from the bone marrow^27^. MPs play an essential role in mouse heart development^24–25,28^. They have been shown to contribute to electrical conductance^28^, valvular remodeling^25^, lymphatic channel development^29^, and endothelial network patterning^24,29^. The role of MPs in human heart development remains unclear, but we hypothesize they play a role because they are present in human embryonic hearts as early as 4-6 gestational weeks^30,31^. To model tissue-resident MPs in the hHOs, we sought to create a hHMA platform. Previously, coculture studies with monocytes have demonstrated that monocyte derivatives model the development, maintenance, pathology, and response to pathology of their respective tissues *in vitro* ^32–36^. Furthermore, *in vitro* coculture experiments with monocytes, MPs, and cardiomyocytes have been used in an attempt to model the pathogenic features of arrhythmias and COVID-19^37–40^. However, these reductionistic systems lack true cardiac cellular complexity and proper anatomical organization. Most recently, MPs, cardiomyocytes and fibroblasts have been incorporated into engineered heart tissue to understand the impact of MPs on the function, cellular, and molecular biology of the tissues^41^.

Atrial fibrillation (AF) is the most common cardiac arrhythmia^42^; characterized by irregular and often rapid heart rhythms, which can lead to severe complications such as stroke and heart failure. Inflammation has been linked to the pathogenesis of AF^43–45^, with particular focus on the NLRP3 inflammasome^46^, a multiprotein complex that plays a crucial role in the innate immune response. Multiple clinical sources point to activation of the NLRP3 inflammasome due to chronic inflammation as a critical step for developing arrhythmogenic disease. In this context, inflammasome triggering is essential in MPs and atrial cardiomyocytes and has been shown to contribute to electrical remodeling, fibrosis, and structural changes in the heart, all of which are hallmarks of AF. As such, NLRP3 inflammasome activation is increasingly recognized as a critical factor in promoting arrhythmogenesis and sustaining AF, making it a potential therapeutic target for managing this condition.

Here, we hypothesized that introducing cardiac MPs in heart organoids would significantly impact organoid development. Using established monocyte differentiation protocols from hPSCs^47–49^ we created a highly reproducible, scalable, and high-content-friendly protocol for hHMA generation, allowing for the investigation of MPs and their impact on human heart development and disease. Using this protocol, we created an *in vitro* atrial arrhythmogenesis model and validated chronic inflammasome activation as a trigger for arrhythmias in human heart tissue.

## RESULTS

### A developmentally inspired strategy to efficiently integrate autologous hPSC-derived embryonic monocytes in hHOs

Our first objective was to develop a protocol that incorporated MPs in hHOs. To ensure hHOs did not contain endogenous MPs, we looked at scRNA-seq data UMAPs of combined 26-day (d) and 15- d hHOs, which showed no gene expression of any classic MP markers, such as *PTPRC* (CD45), *CD68*, *CD163*, or *CD14* (Figure S1A). We also could not detect any endocardial-derived MPs^50,51^ although we found numerous *NFATC1^+^* / *CDH5^+^* cells typically associated with the endocardium in the hHOs (Figure S1A), as described previously^17^. After verifying that there were no pre-existing MPs in hHOs, we tested if autologous hPSC-derived monocytes would differentiate into cardiac tissue-resident MPs if added to hHOs with the right timing. To estimate this timing, we relied on MP integration events observed in the developing mouse^24,29,52^ and human heart^53,54^. In the mouse, the first heartbeats occur at E8.5^55^, and yolk sac-derived MPs don’t appear until E10.5^51^. The epicardium seems to be required for cardiac seeding by the yolk sac-derived MPs^56^, thus, the addition of monocytes to the hHOs must occur after the day 7 CHIR99201 exposure, which induces the formation of the pro-epicardial organ in the hHO model^17^. Coronary vasculature in the mouse heart forms at E13.5^24^, in hHOs early vasculature is detected as early as day 12 of hHO differentiation^17^. Thus, we determined that the first monocyte addition should occur between day 7 and day 12 of hHO differentiation. hHOs^17^ also exhibited increased expression of genes related to leukocyte adhesion, monocyte differentiation, and positive regulation of MP chemotaxis around day 7 of differentiation (Figure 1A), supporting this time window for monocyte integration. We also noted that VCAM1 expression, an adhesion molecule necessary for cardiac-macrophage interaction and recruitment, was high at day 11 and colocalized with TNNT2 (Figure 1B), further suggesting cardiac cells were primed, a finding previously reported in embryonic mouse hearts^57^. Several studies have demonstrated it takes approximately four days for monocytes to differentiate into MPs *in vitro*^47,48^. Considering all these observations, we reasoned that monocyte additions should start on day 8 of differentiation.

**Figure 1.**
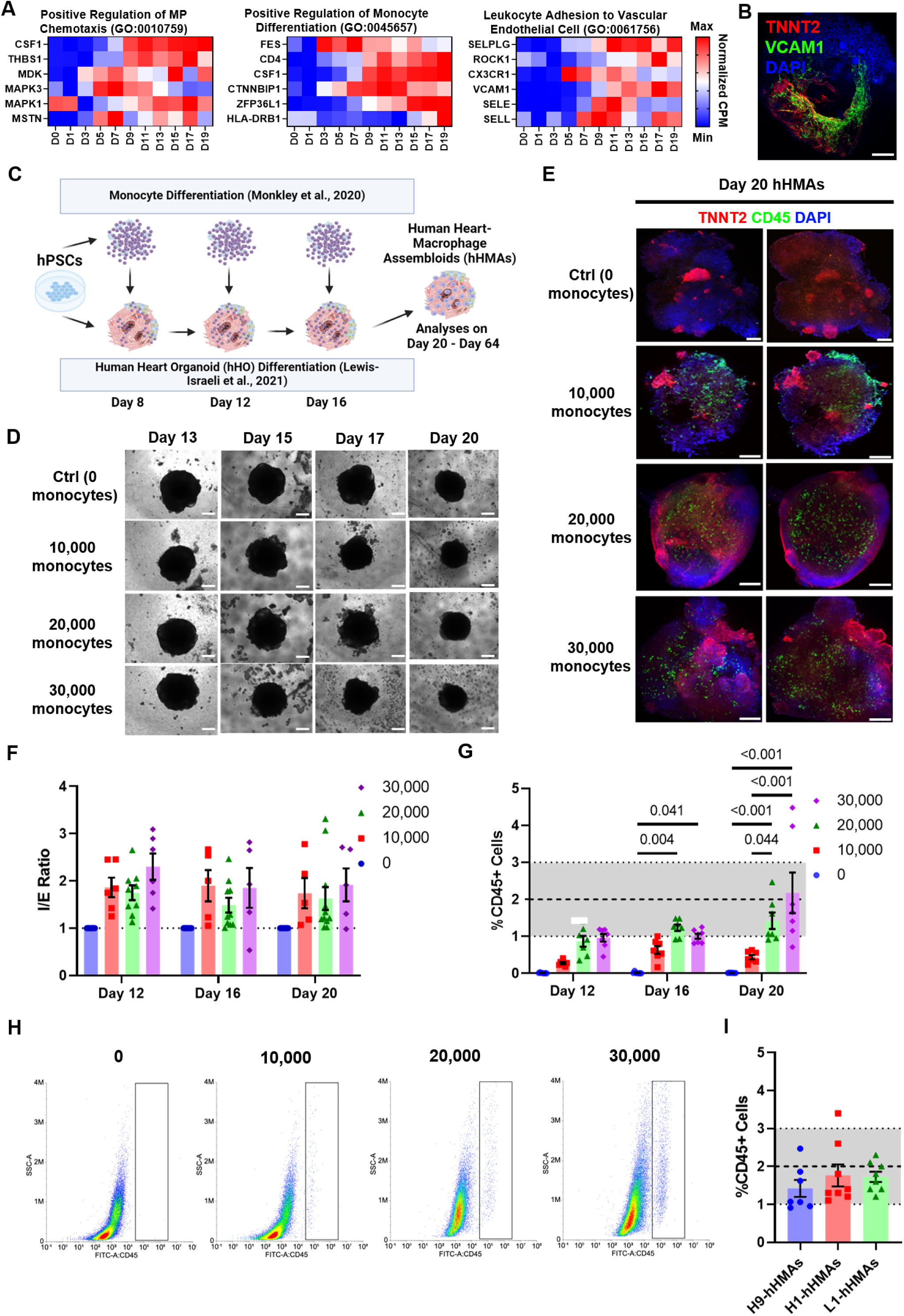
Autologous hPSC-derived embryonic monocytes efficiently integrate into human heart organoids. (A) Heat map of normalized expression of genes involved in Leukocyte Adhesion to Vascular Endothelial Cell, Positive Regulation of Monocyte Differentiation, and Positive Regulation of Macrophage of Chemotaxis in hHOs from bulk RNA-seq data from Day 0 to Day 19 of hHO differentiation. (B) High magnification confocal IF image for DAPI (blue), VCAM1 (green), and TNNT2 (red), indicated by the white box in a. Scale = 200 µm. n = 8. (C) Flowchart depicting the hHMA generation protocol. (D) Phase contrast microscopy images depicting hHMAs at day 13, 15, 17, and 20 with incremental additions of monocytes (10,000, 20,000, and 30,000) at day 8, 12, and 16. After each monocyte addition, the hHO and monocytes were subject to two pipet resuspensions. Scale = 200 µm. n = 8. (E) Representative confocal immunofluorescence images displaying the exterior (left panel) and interior (right panel) of day 20 hHMAs stained for DAPI (blue), CD45 (green) and TNNT2 (red). Top row Scale = 200 µm. n ≥ 6. (F) Quantification of immune cell (CD45^+^) integration using z-planes from immunofluorescent images depicting the interior (three images per hHMA) and exterior of individual hHMAs with incremental monocyte additions of 0 (blue), 10,000 (red), 20,000 (green), and 30,000 (purple) monocytes, represented as an I/E ratio: I/E Ratio = (%CD45^+^ cells located on the interior of the hHMAs/%CD45^+^ cells located on the exterior of the hHMAs). n ≥ 5. (G) Cell quantification of immune cells using flow cytometry data for CD45^+^ cells as a percentage of total cells at day 12, 16, and 20 with incremental monocyte additions of 0 (blue), 10,000 (red), 20,000 (green), and 30,000 (purple) monocytes (n ≥ 7). The gray shaded area indicates the expected physiologic range of macrophages in the developing human heart (1-4%). n ≥ 5. (H) Representative flow cytometry data showed day 20 hHMAs (left to right) with incremental monocyte additions of 0, 10,000, 20,000, and 30,000 cells. The black box represents a gate for CD45^+^ cells. (I) Quantification of % CD45^+^ cells as determined by flow cytometry in three cell lines: H9-hESCs, H1-hESCs, and L1-hiPSCs. The gray shaded area indicates the typical physiologic range of macrophages in the developing human heart (1-3%). n ≥ 7. For all graphs: Value = mean ± s.e.m., 1-way ANOVA multiple comparison test.

Monocytes were derived using a previously established differentiation protocol^49^ (Figure S1B) and were purified using magnetic-antibody cell sorting (MACS) for CD14, since >94% of human cardiac tissue-resident MPs express CD14^58^. CD14^+^ isolated monocytes also expressed PTPRC (CD45), CD14, CD68, and CD163 (Figures S1C-S1G), classical monocyte markers. Monocyte additions started on day 8, with follow-up additions on days 12 and 16 to mimic sustained MP seeding during development (Figure 1C). Monocytes were delivered into the standard hHO-containing medium in organoid plates to promote contact and resuspended twice with each addition. No significant differences in hHO morphology were noted early on after monocyte additions of 10,000, 20,000, and 30,000 cells per day (Figure 1D), but a dose-dependent incorporation of CD45+ cells was noted (Figures 1E and S1H). Z-stack immunofluorescent confocal microscopy images showed a preference for CD45^+^ cells to populate the interior compartments of the organoids versus the hHO exterior (Figure 1E, F). By day 20 of hHO differentiation, adding 20,000–30,000 monocytes on days 8, 12, and 16 achieved physiologically relevant CD45^+^ populations (∼1-3% of total cells) in hHOs, consistent with scRNA-seq data from human embryonic heart scRNA-seq datasets^59^ (Figures 1G-H and S1I). We differentiated monocytes and hHOs from three different hPSC lines and observed similar results for monocyte integration, as determined by CD45^+^ positive content in hHOs (Figure 1G). Thus, we concluded that hHOs can be successfully seeded and integrated with autologous MPs. We called this new organoid model human heart-macrophage assembloids (hHMAs) since the system fits the assembloid definition of being generated by putting in contact with two different organ tissue types followed by developmental self-organization-driven integration.

### Integrated monocytes adopt embryonic cardiac tissue-resident MP fates and persist in hHMAs over time

To verify that the added monocytes did become cardiac tissue-resident MPs, we used immunofluorescence microscopy to validate colocalization of CD45, a general hematopoietic cell marker, with CD163, a monocyte-MP marker (Figure 2A). Gene expression analysis by qRT-PCR showed a significant level of genes expressed by embryonic cardiac tissue-resident MPs in day 20 hHMAs vs non-integrated hHOs (Figure 2B). scRNA-seq UMAPs of hHMAs showed specific expression of embryonic tissue-resident MP markers in the expected assigned MP cluster (Figure 2C). We also noted the lack of *CCR2 expression*, a chemokine receptor not expressed by embryonic-derived tissue-resident MPs in the heart^27^. Tissue-resident MPs seed the heart early in embryonic development and persist in the heart for long periods into adulthood, significantly supported by endogenously-produced factors produced by cardiac cells^26,60,61^. To test the longevity of the MPs in hHMAs, we cultured hHMAs until day 60. We noticed that hHMAs morphologically changed over time compared to hHOs (Figures 2D-F). While the area of the organoids did not significantly change, hHMAs had significantly higher circularity indexes than hHOs. hHMAs also presented less organoid-associated debris. Immunofluorescence microscopy studies confirmed the presence of MPs through day 60 of hHMA culture (Figures 2G-I). MP numbers peaked at day 28 and then gradually decreased ∼25% by day 36, remaining stable up to day 60 (Figure 2H), but always stayed within physiologically relevant levels (1-3% of the total cardiac cell population). MPs did not affect beating in the hHMAs (Figure S2A). Moreover, from day 15 to day 26, MPs had decreased expression of genes associated with M1 or M2 MPs, *CCL2*^62^, and MIF50 (Figure S2B), confirming their tissue-resident MP fate. Gene ontology of differentially expressed genes between MPs at day 26 hHMAs vs. day 15 hHMAs showed significantly higher levels of genes encoding enzymes for lipid and cholesterol metabolism (Figure S2C), suggesting metabolic maturation, a process associated with tissue-resident MPs^63^.

**Figure 2.**
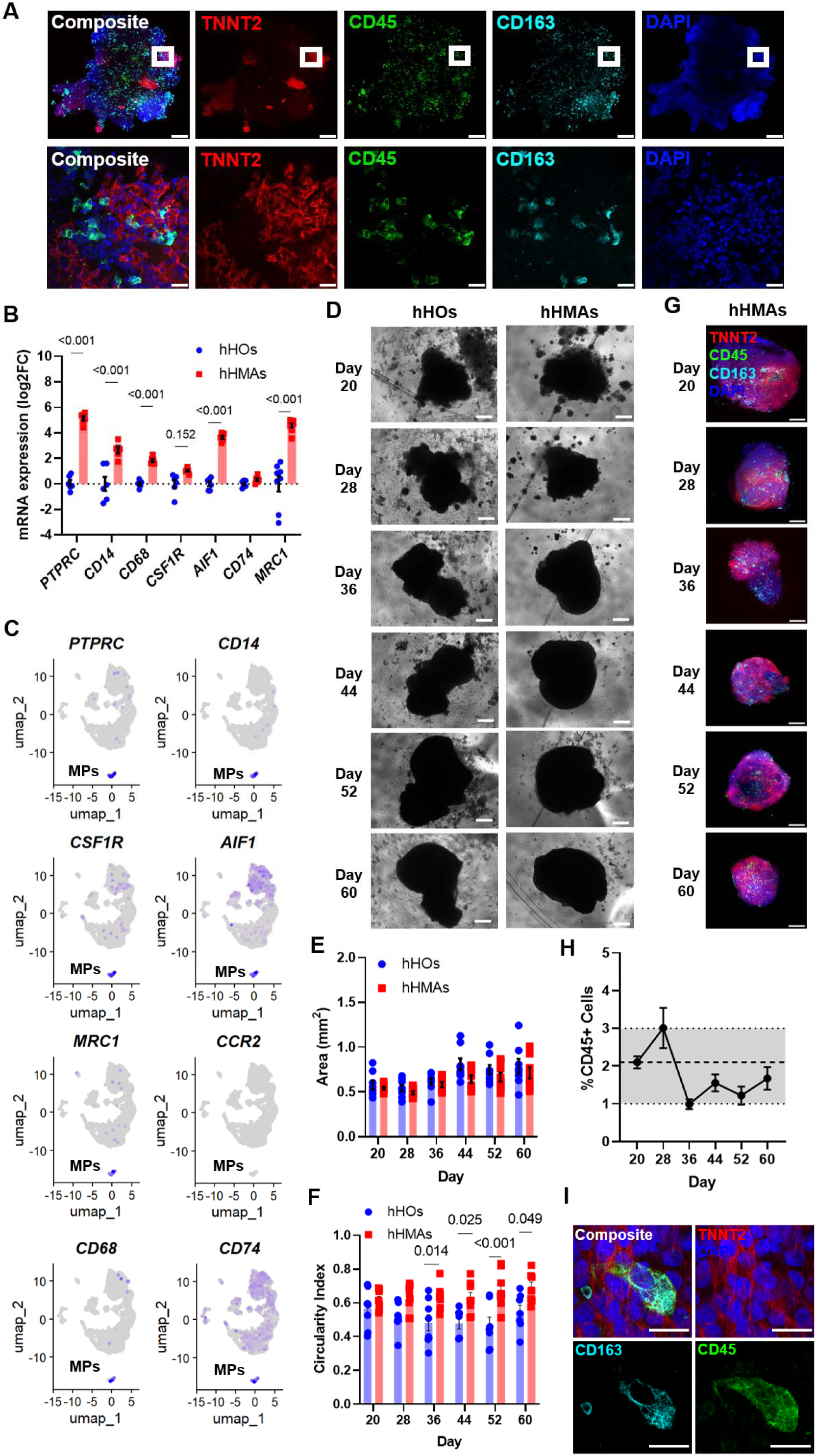
Integrated monocytes acquire embryonic cardiac tissue-resident MP fate and persist long-term. (A) Representative confocal immunofluorescence images for DAPI (blue), CD45 (green), CD163 (cyan), and TNNT2 (red), in day 20 hHMAs at low (top row) and high magnification (bottom row). The white box indicates the location where high-magnification images were taken from the low-magnification image. n = 8. Top row scale = 200µm. Bottom row scale = 20µm. (B) RT-qPCR analyses on day 20 hHMAs for classical proteins markers expressed in cardiac embryonic macrophages, PTPRC (CD45), CD14, CD68, CSF1R, AIF1, CD74, MRC1 (CD206), and CCR2. n ≥ 6. Value = mean ± s.e.m., Two-way ANOVA with multiple comparison test. (C) UMAP feature plots displaying relative expression for cardiac embryonic tissue-resident macrophages (MPs) markers, PTPRC (CD45), CD14, CD68, MRC1, CSF1R, AIF1, and CD74 in hHMAs (day 15 and day 26 hHMA overlay). CCR2 is included and is not classically expressed in cardiac embryonic tissue-resident MPs. Color intensity represents the relative value of gene expression per gene. The MP cluster is labeled on each UMAP plot. (D) Phase-contrast microscopy images depicting individual control hHOs and hHMAs at day 20, 28, 36, 44, 52, and 60. hHMAs had 20,000 monocytes added to them on days 8, 12, and 16. Scale = 200µm. (E) Quantification of control hHOs (blue) and hHMAs (red) area under phase-contrast microscopy at day 20, 28, 36, 44, 52, and 60. n = 8. Value = mean ± s.e.m. One-way ANOVA with Turkey’s correction with multiple comparisons. (F) Quantification of control hHOs (blue) and hHMAs (red) circularity under phase-contrast microscopy at days 20, 28, 36, 44, 52, and 60. n = 8. For all graphs: Value = mean ± s.e.m. One-way ANOVA with Turkey’s correction with multiple comparisons. (G) Representative IF images of day 20, 28, 36, 44, 52, and 60 hHMAs stained for TNNT2 (red), CD45 (green), CD163 (cyan), and DAPI (blue). n = 8. Scale = 200µm. (H) Cell quantification of CD45+ cells using flow cytometry for CD45+ cells as a percentage of total cells in day 20, 28, 36, 44, 52, and 60 hHMAs. The dashed line represents the MP population on day 20. The gray shaded area indicates the expected physiologic range of macrophages in the developing human heart (1-4%). n ≥ 7. Value = mean ± s.e.m. (I) High magnification IF images of a day 60 hHMAs stained for TNNT2 (red), CD45 (green), CD163 (cyan), and DAPI (blue). n = 8. Scale = 20µm.

To study the structure of day 26 hHMAs, we employed optical coherence tomography (OCT) on day 26 hHOs and hHMAs (Figure S2D-J; Videos S1 and S2). OCT imaging revealed significantly larger interconnected hypodense regions within hHMAs not present in hHOs, suggesting MPs contributed to chamber morphogenesis and maturation (Figure S2F). OCT imaging also showed that hHMAs were considerably more compact than hHOs, without altering the number of chambers within the organoids, suggesting improved cardiac muscle compaction (Figure S2G). This data confirms that integrated monocytes acquire embryonic cardiac tissue-resident MPs fates, persist in hHMAs without exogenous cytokine additions, and alter the structure of the native organoids, suggesting developmental impact.

### Single-cell transcriptomics of hHMAs reveals the effect of tissue-resident MPs on cardiac development

scRNA-seq was performed on day 15 hHOs, day 15 hHMAs, day 26 hHOs, and day 26 hHMAs to determine how MP integration altered organoid cell populations and gene expression (Figures 3 and S3). We made cluster assignments for each scRNA-seq sample based on the top 5 differentially expressed genes per cluster (Figure S3A). MPs constituted 1.7% of all cells on day 15 and 2.1% of all cells on day 26 hHMAs (Figures 3A-B and S3B-C), comparable to embryonic human hearts^30^. Furthermore, significant cluster population changes were noted with MP integration (Figures 3B-C and 3SB-C). In day 26 hHMAs, cardiac fibroblasts (CFs), ventricular cardiomyocytes (VCMs), and valvular cells (VCs) had significantly larger populations, while conductance cells (CCs), cardiac progenitor cells (CPCs), endothelial cells (ENCs), epicardial cells (EPCs), stromal cells (SCs), and proliferating pro-epicardial derived cells (PEDCs) had significantly smaller populations as compared to day 26 hHOs. Integrated scRNA-seq datasets from previously published human embryonic hearts revealed that hHMAs were remarkably similar and that all the key cell populations found in age-matched embryonic human hearts were present in the hHMAs (Figures 3D and 3SD)^30,64^. Dot plot analyses further demonstrated that cluster assignments appropriately expressed genes commonly associated with each cluster (Figure 3E), MPs specifically expressed *PTPRC*, *CD14*, *CD163*, *CSF1R*, and *CD68,* all of which are cardiac tissue-resident MP markers. We also performed gene set enrichment analyses (GSEA) to show that hHMAs significantly expressed gene sets associated with tissue-resident MPs function, such as the innate immune response, leukocyte migration, and IL-10 signaling (Figures 3F-G and 3SE).

**Figure 3.**
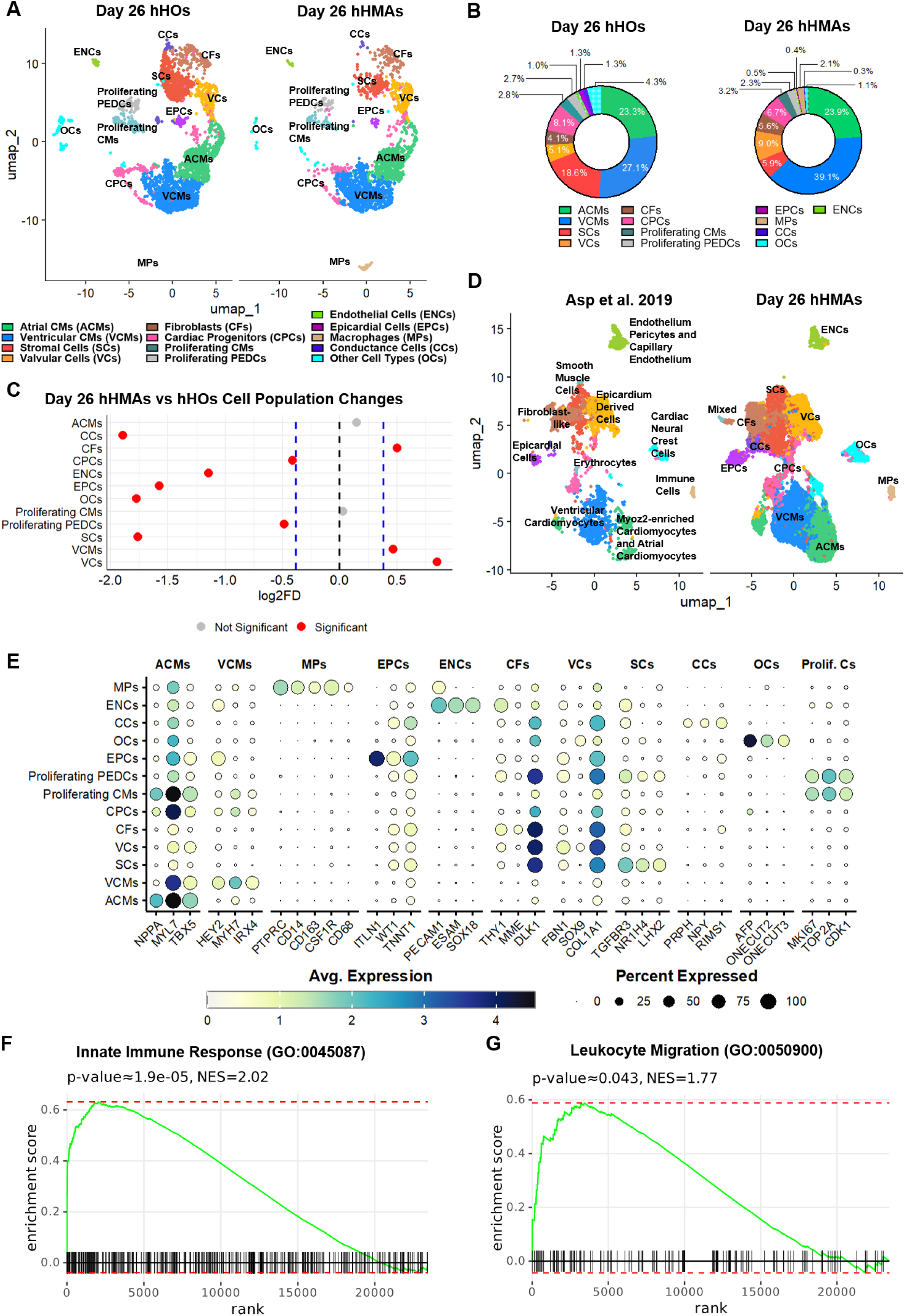
Cardiac tissue-resident MPs reshape the cellular composition of heart organoids and increase developmental fidelity. (A) UMAP dimensional reduction plots of integrated scRNAseq data for each condition (from left to right): Day 26 hHOs and Day 26 hHMAs. Cluster identity labels are on the UMAP plots. (B) Quantification of total cell count percentages per cluster for Day 26 hHOs and Day 26 hHMAs. Colors of regions correspond to the adjacent legend. (C) Log_2_ fold differences (log_2_FD) in the proportion of cells across clusters between Day 26 hHOs and Day 26 hHMAs. Clusters highlighted in red indicate an average |log2 fold difference| greater than 0.38 compared to hHOs (permutation test; n = 10,000). (D) UMAP dimensional reduction plots of integrated scRNAseq data of embryonic human hearts from Asp et al. 2019 (left) and Day 26 hHMAs (right). Cluster labels from Asp et al. 2019 are preserved from the original text (left) and hHMA labels from Figure 2A (right). (E) Dot plot of differentially expressed genes in each cluster for each condition. The color indicates the average expression level across all cells, and the circle’s size means the percentage of cells within a particular cluster that expresses the respective gene. (F-G) Gene set enrichment plots of differential gene expression between hHMAs and hHOs for Innate Immune Response Gene Ontology 2023 (F) and Leukocyte Migration Gene Ontology 2023 (G).

### Tissue-resident MPs program cell-cell communication in hHMAs

One of the advantages of our model is the effective control of the organoid environment, allowing us to perform *in vitro* cell communication studies in organoids and through the collection of cell culture media without interference from other organs or other difficult-to-control exogenous factors^65^. We used scRNA-seq ligand-receptor (L-R) and differential L-R (Diff L-R) analysis to identify the production and interaction of macrophage-associated cytokines with cardiac cells. We noted that MPs had a disproportionately large number of interactions with every other cardiac cell type in hHMAs (Figure 4A). Interestingly, of the top 20 most prevalent L-R interactions from MPs in hHMAs, 7 involved *SPP1* (osteocalcin)(Figure 4B), which is also expressed by MPs in human embryonic hearts^30,64^, suggesting a potentially unexplored critical role for SPP1 in heart development. Diff L-R interactions significantly upregulated in non-MP clusters were genes associated with regulating the integrin-mediated signaling pathway, cell-matrix adhesion, and molecular processes related to proteoglycan binding (Figure 4C and S4A-B). Diff L-R analyses between day 26 hHMAs and hHOs found that tissue-resident MPs upregulate interactions from CFs, VCs, proliferating PEDCs, and SCs while decreasing cell signaling from VCMs (Figure S4C). Overall, MP inclusion resulted in the global upregulation of L-R interactions compared to control hHOs (Figure S4D). Gene ontology analyses on the most represented ligands produced by MPs showed contributions to epithelial-to-mesenchymal transition, the complement cascade, angiogenesis, the inflammatory response, and myogenesis (Figure 4D). Furthermore, we confirmed that the top 15 represented MP ligands were expressed more in day 15 and 26 hHMAs compared to hHOs (Figures S4E).

**Figure 4.**
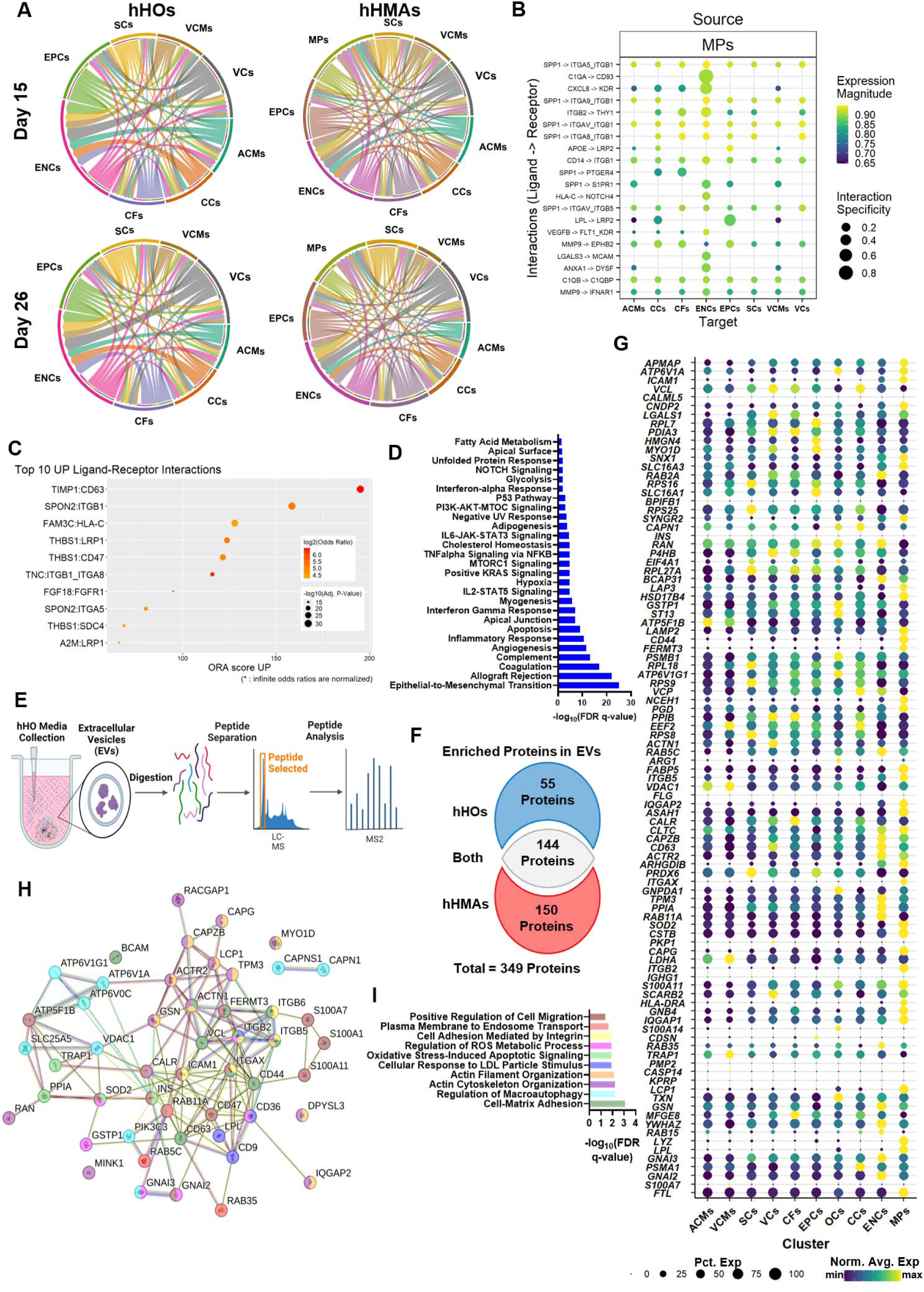
Cardiac tissue-resident MPs program hHMA heart development by activating unique signaling pathways. (A) Circularized Ligand-Receptor plots for day 15 hHOs, day 26 hHOs, day 15 hHMAs, and day 26 hHMAs representing the top 10 ligand-receptor interactions for each specific cell type. (B) Ligand-Receptor dot plot displaying the top 20 ligand-receptor interactions between MPs and specific cell types in day 26 hHMAs. (C) Top 10 upregulated ligand-receptor interactions from the differential ligand-receptor analysis between day 26 hHMAs vs day 26 hHOs. (D) Hallmark gene ontology biological processes that MP ligands contribute to in hHMAs. (E) Schematic of EV isolation and proteomic analysis from hHOs and hHMAs. (F) Venn diagram displaying the distribution of proteins found in EVs unique to hHO culture (blue), hHMA culture (red), and both cultures (light grey). (G) Dot plot displaying differential expression of the most prevalent proteins in EVs from only hHMAs by cluster. Color is indicative of the normalized average gene expression between each cluster, and the size of the circle is indicative of the percentage of cells within the cluster that express the respective gene. (H) String plot of proteins found in hHMA EVs involved in the GO Biological Processes in I. Colors correlate to the color of bars found in I. (I) 10 significantly upregulated GO Biological Processes using the proteins found in hHMA EVs.

EVs are a newly recognized cellular communication mechanism, and macrophages are known for EV release and cardiovascular physiology^66^. We performed LC-MS proteomic analyses of the EVs present in the cell culture medium of day 20-26 hHMAs and compared these to media from hHOs at the same timepoints (Figure 4E). hHMA EVs showed 150 unique proteins, compared to 55 unique proteins in hHOs (Figure 4F). Many unique proteins were expressed predominantly by MPs, such as LYZ, LPL, CD44, FERMT3, FTL, HLA-DRA, ITGB2, ITGAX, CAPG, and SOD2. In contrast, other proteins were expressed by different types of cells but exclusively in the presence of MPs, such as CALR, LDHA, TRAP1, MFGE8, PRDX6, VCP, RAN, and VCL (Figure 4G), suggesting MP-directed programming of cardiac cells is necessary for their production. Proteins found in the EVs of both hHMAs and hHOs were predominantly housekeeping cytoskeletal proteins (e.g. keratins) (Figure S4F, G). Gene ontology and STRING analyses of EV-proteins in hHMAs showed increased activation of endosomal transport, cellular response to LDL, integrin binding, ATP synthase activity, cell-matrix-adhesion, regulation of macroautophagy, actin-filament organization, and positive regulation of cell migration (Figures 4H and 4I). Whereas gene ontology and STRING analyses of the 20 most represented unique EV- proteins in hHOs were related to system development, regulation of cell growth, and regulation of cell population proliferation (Figures S4H-K). In summary, these data show EVs from MPs significantly contribute to cell-to-cell signaling and communication within the developing human heart and identify new proteins and genes of interest for future developmental studies.

### MPs functionally remodel hHMAs by promoting efferocytosis, ECM organization, sarcomere growth, and ventricular morphogenesis

We wanted to determine if MPs exhibit preferential localization in certain parts of the hHMAs. Immunofluorescence staining and quantification of day 20 hHMAs in low-magnification confocal microscopy images showed MP predominantly localized to TNNT2+ areas, WT1+ areas, and chamber regions of the organoids (Figure S5A, B). WT1 is expressed by VCs, CFs, SCs, and EPCs in hHMAs (data not shown), indicating MPs accumulate more with cardiac interstitial cells than cardiomyocytes. A recent study showed MPs enhanced the contractility and morphogenesis of VCMs from hPSC-derived engineered heart tissues^41^. Combined with our findings of a higher composition of VCMs in day 26 hHMAs (Figure 2A-C), we sought to find evidence that MPs enhance hHMA contractility and ventricular cardiomyocyte morphogenesis through transcriptomics and immunofluorescence microscopy. Differential gene expression showed gene set enrichment for cardiac muscle contraction in hHMAs vs. hHOs (Figure 5A). Immunofluorescence microscopy of day 26 hHMAs showed significantly higher contractile protein expression (TNNT2^+^, MYL3^+^) than hHOs (Figures 5B-C and S5C-D). High-magnification confocal images of sarcomeres in day 26 hHMAs revealed longer sarcomere structures compared to hHOs (Figure 5D and 5E), a sign of cardiomyocyte maturity^67^. Genes involved in sarcomere organization, cardiac ventricle morphogenesis, and ventricular muscle tissue development were elevated in hHMAs at day 15 and day 26 compared to hHOs (Figure 5F). Furthermore, MPs are known to participate in homeostatic functions like efferocytosis^68^ and cardiomyocyte exopher removal^69^ in the heart. GSEA for differentially expressed genes between hHMAs and hHOs indicated significant enrichment in phagocytosis recognition pathways (Figure 5G), highlighting the involvement of MPs in clearing apoptotic cells and cellular debris within hHMAs. Phase-contrast microscopy of hHMAs and hHOs over a time course from day 20 to day 60 showed significantly less cellular debris (Figure S5E and S5F), which was defined as the shadow surrounding the organoids from phase-contrast images. This is supported by pseudo-bulk transcriptomic data that indicated upregulated gene expression for processes associated with phagocytosis, apoptotic cell clearance, and ECM remodeling in hHMAs, particularly at day 26, compared to hHOs (Figure S5G). CALR is an “eat-me” signal expressed by both live and apoptotic cells that is recognized by MPs^70,71^. Confocal imaging of day 26 hHMAs displayed localization of CD45+ MPs with CALR+ cells (Figure 5H-K), indicating physiological efferocytotic function. Quantitative analyses revealed that a significant proportion of CD45+ MPs were associated with CALR+ regions. Immunofluorescence staining for MERTK, a receptor involved in cardiac MP efferocytosis^72,73^ and disposal of cardiac exophers^69^, showed significantly increased MERTK^+^ puncta in CD45+ MPs compared to other cell types within day 26 hHMAs. These findings underscore the role of MPs in facilitating efferocytosis within hHMA culture, a function traditionally associated with MPs in cardiac tissue.

**Figure 5.**
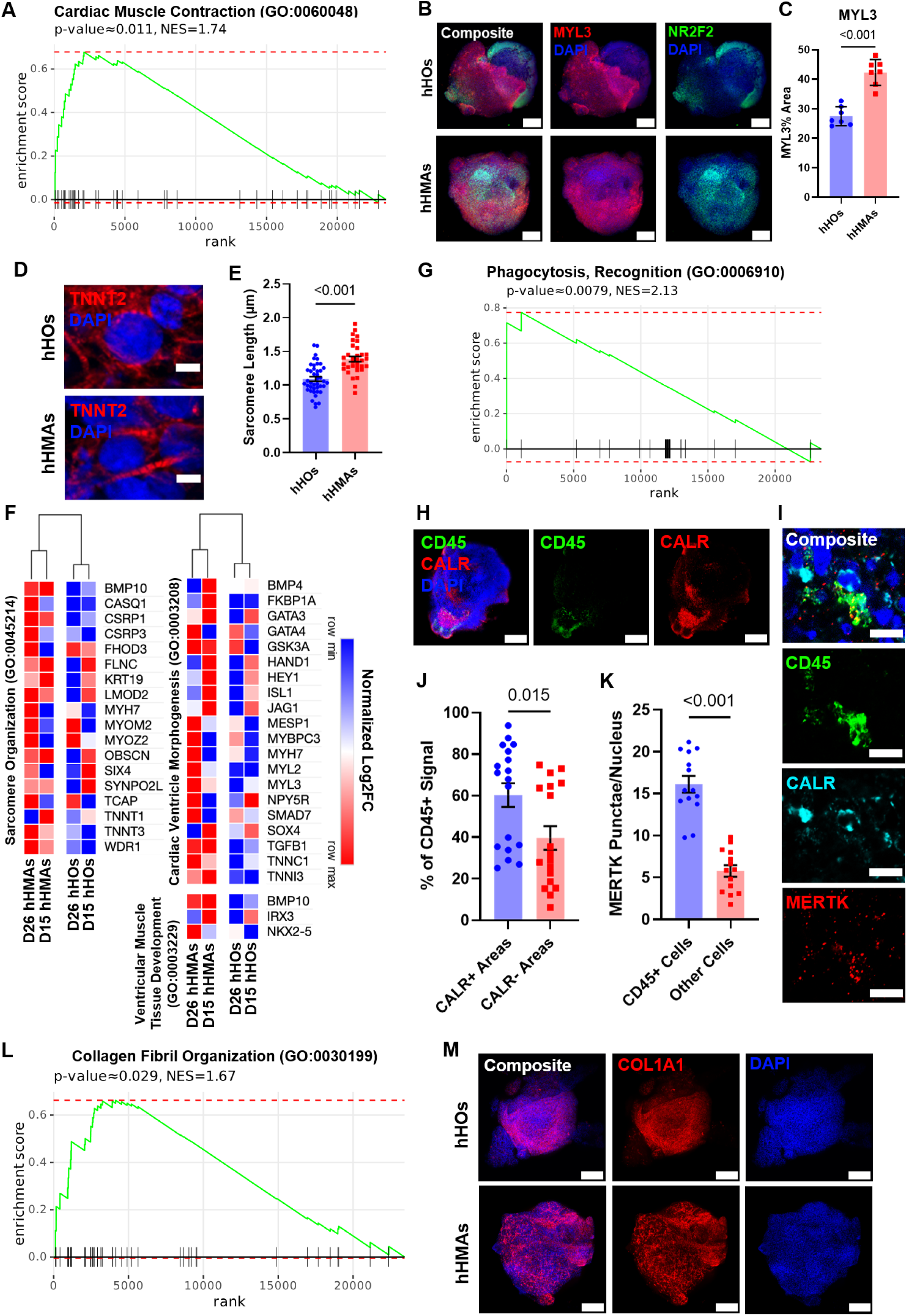
Autologous cardiac tissue-resident MPs modify the extracellular matrix, promote efferocytosis, sarcomere growth and ventricular morphogenesis. (A) Gene set enrichment plots of differential gene expression between hHMAs and hHOs for Cardiac Muscle Contraction, GO Biological Process 2023. (B) Representative low-magnification confocal IF image for DAPI (blue), MYL3 (red), and NR2F2 (green) in hHOs (top row) and hHMAs (bottom row). n = 7. Scale Bar = 200µm. (C) Quantification of MYL3+ area as a percentage of total area in confocal IF images of day 26 hHOs and hHMAs averaged across 5 z-slices per organoid. n = 7. Value = mean ± s.e.m., Student’s t-test. (D) Representative high-magnification confocal IF images of sarcomeres in day 26 hHOs (left) and hHMAs (right). n ≥ 11. Scale = 5µm. (E) Quantification of sarcomere length, measured from z-line to z-line between TNNT2+ signal, from high-magnification confocal IF images of sarcomeres in hHOs and hHMAs. n_organoids/sarcomeres_≥ 11/33. Value = mean ± s.e.m., Student’s t-test. (F) Heatmap depicting normalized log2 fold change for genes related sarcomere organization, cardiac ventricle morphogenesis, and ventricular muscle tissue development in day 15 hHOs, day 26 hHOs, day 15 hHMAs, and day 26 hHMAs. Red correlates to maximum relative expression and blue correlates to minimum relative expression. (G) Gene set enrichment plots of differential gene expression between hHMAs and hHOs for Phagocytosis, Recognition, GO Biological Process 2023. (H) Representative confocal IF images of Day 26 hHMAs for DAPI (blue), CD45 (green), and CALR (red). n = 18 organoids. Scale Bar = 200µm. (I) Representative confocal IF images of Day 26 hHMAs for DAPI (blue), CD45 (green), MERTK (red), and CALR (cyan). n = 14. Scale Bar = 10µm. (J) Quantification of CD45+ signal in CALR+/- regions of interest in Day 26 hHMAs. n = 18 organoids. For graph: Value = mean ± s.e.m., Student’s t-test. (K) Quantification of MERTK+ punctae in CD45 cells and all other cells in Day 26 hHMAs. n = 14 organoids. For graph: Value = mean ± s.e.m., Student’s t-test. (L) Gene set enrichment plots of differential gene expression between hHMAs and hHOs for Collagen Fibril Organization, GO Biological Process 2023. (M) Representative confocal IF image for DAPI (blue) and COL1A1 (red), displaying the exterior of day 26 hHMAs at low magnification. n ≥ 7. Scale Bar = 200µm.

MPs are also known to interact with, and remodel, the ECM^74^. GSEA identified significant upregulation of genes involved in collagen fibril organization from hHMAs compared to hHOs (Figure 5L), suggesting MP- driven ECM remodeling. Confocal images of day 26 hHMAs stained for COL1A1 display altered collagen deposition in the MP-containing regions (Figure 5M). Pseudo-bulk transcriptomic data also shows increased relative expression in genes related to ECM assembly, cardiac epithelial-to-mesenchymal transition, and collagen fibril organization in hHMAs versus hHOs (Figure 5G). RT-qPCR analyses further corroborated these findings by showing increased expression of COL1A1 in day 20 hHMAs compared to hHOs (Figure S5H). Together, these findings suggest that autologous cardiac MPs alter the ECM within hHMAs.

### MPs enhance catabolic processes in hHMAs

GSEA of differentially expressed genes between hHMAs and hHOs identified significant upregulation of glycolytic processes in hHMAs (Figure S6A). A further breakdown of the top 10 gene ontology processes from the 250 most differentially expressed genes in day 26 hHMAs compared to hHOs shows significant enrichment in various catabolic processes, including glycolysis, mitochondrial function, and fatty acid metabolism (Figure S6B). These findings indicate that MPs are essential in modulating metabolic activity in hHMAs. Feature plots of VCMs on day 26 hHMAs and hHOs show increased expression of IGFBP2, IGF2, PGK1, ENO1, and GPI (Figures 6C and 6D). A heatmap illustrating the normalized log_2_ fold change (log_2_FC) of crucial metabolic genes revealed upregulation of glycolysis, insulin-like growth factor receptor (IGFR) signaling, mitochondrial electron transport (cytochrome C to oxygen), fatty acid beta-oxidation, lactate metabolism, and intracellular oxygen homeostasis in hHMAs (Figure S6E). These findings demonstrated that MPs in hHMAs enhance catabolic processes, including glycolysis and mitochondrial function, contributing to the overall metabolic regulation and energy homeostasis in hHMAs.

**Figure 6.**
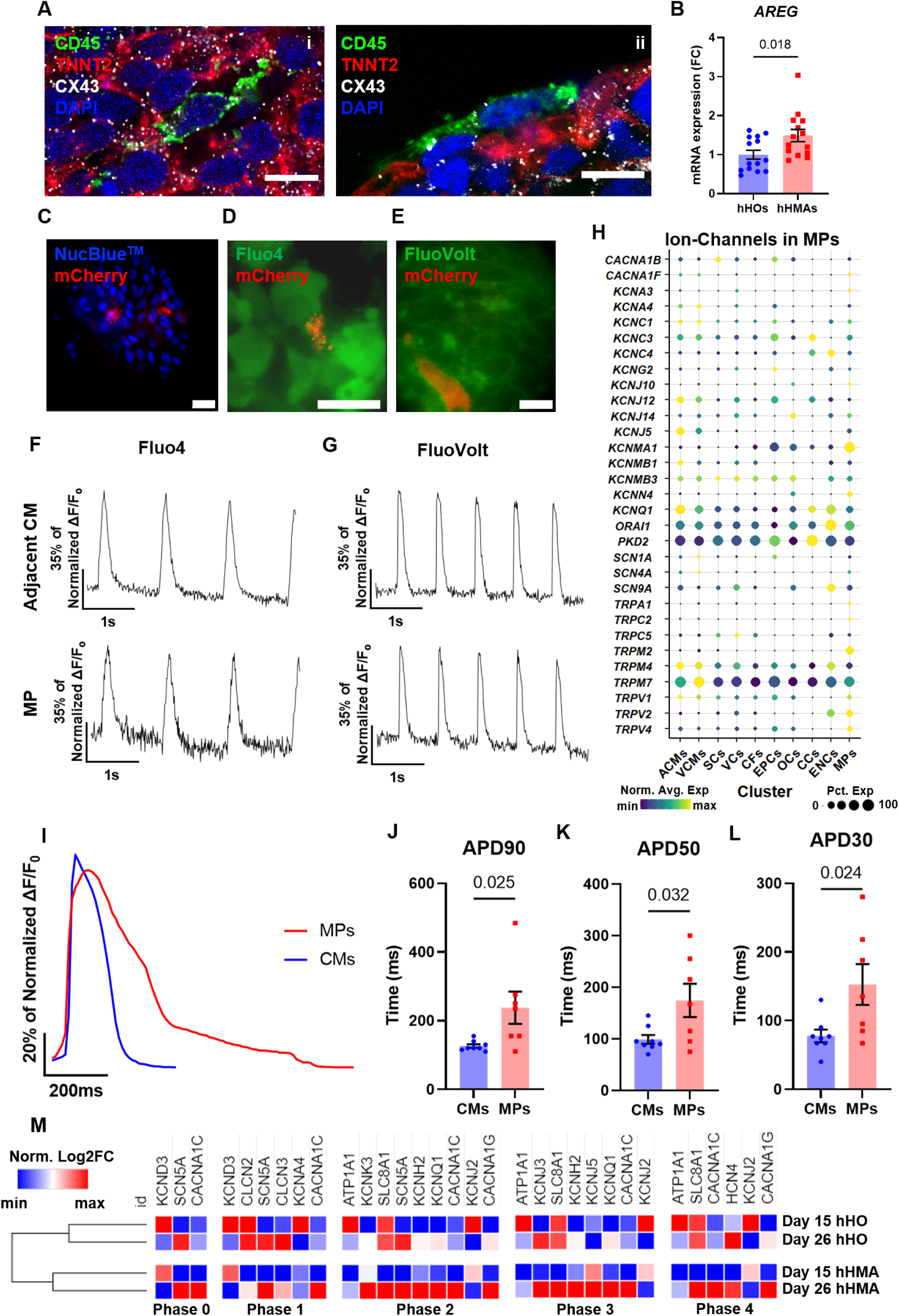
MPs form GAP junctions and contribute to ion signaling in hHMAs. (A) Representative high magnification confocal IF images (i and ii) of MPs for DAPI (blue), CD45 (green), TNNT2 (red), and Cx43 (white) in day 20 hHMAs. N = 10. Scale = 10µm. (B) RT-qPCR analyses on day 20 hHMAs for AREG, a protein expressed by cardiac embryonic MPs when Cx43 GAP junctions are formed. n ≥ 13. Value = mean ± s.e.m., Student’s t-test. (C) Live-cell confocal IF imaging for NucBlueTM (blue) and mCherry (red) positive MPs. n ≥ 39. Scale = 20µm. (D) Representative live-cell confocal IF imaging for Fluo4 dye (green) and mCherry (red) positive MPs. n ≥ 8. Scale = 20µm. (E) Representative live-cell confocal IF image for FluoVolt dye (green) and mCherry (red) positive MPs. n ≥ 8. Scale = 10 µm. (F) Ca2+ transients measured using Fluo-4 live-cell confocal imaging of representative hHMAs in an mCherry-MP (bottom) and an adjacent cardiomyocyte (top). n ≥ 8 (G) Action potentials were measured using FluoVolt live-cell confocal imaging of representative hHMAs in an mCherry-MP (bottom) and an adjacent CM (top). n ≥ 8. (H) Dot plot displaying differential gene expression of ion channels in at least 0.01% of MPs with an average normalized expression of 0.1. Color is indicative of the normalized average gene expression between each cluster and the size of the circle is indicative of the percentage of cells within the cluster that express the respective gene. (I) Average action potential waveforms of CMs and mCherry-MPs captured by Fluovolt live-cell imaging. n ≥ 7. (J – L) Quantified APD90 (J), APD50 (K), and APD30 (L) of CMs and mCherry-MPs captured by Fluovolt live- cell imaging. n ≥ 7. Value = mean ± s.e.m., Student’s t-test. (M) Heatmap depicting normalized log2 fold change for genes involved in Phase 0, 1, 2, 3, and 4 of the cardiac action potential in day 15 hHOs, day 26 hHOs, day 15 hHMAs, and day 26 hHMAs. Red correlates to maximum relative expression and blue correlates to minimum relative expression.

### MPs form GAP junctions with cardiomyocytes and contribute to the electrophysiology profile of hHMAs

Studies in the last decade have highlighted the contribution of MPs to cardiac electrophysiology^28^. We wanted to determine if MPs participated in the electrophysiology of hHMAs. High-magnification confocal immunofluorescence images of day 20 hHMAs stained for CD45, TNNT2, and connexin 43 (CX43) showed the formation of GAP junctions between MPs and cardiomyocytes (Figure 6A). CX43 expression in the regions of MP-cardiomyocyte contact suggested functional coupling between the two cell types. Analysis of bulk RNA-seq from day 20 hHMAs and hHOs revealed a significant upregulation of AREG (Figure 6B), a protein associated with the formation of CX43-mediated GAP junctions between MPs and cardiomyocytes in mice hearts^75^, indicating that MPs in hHMAs form GAP junctions with cardiomyocytes. Live-cell confocal imaging of genetically labeled mCherry MPs (Figure 6C-G; Video S3) showed live MPs integrated in hHMAs, and subsequent functional imaging with Fluo-4 dye (for Ca2+ activity) and FluoVolt dye (for membrane potential) illustrated MPs with synchronized calcium signaling and action potentials alongside adjacent cardiomyocytes. To elucidate which MP-specific ion channels may be contributing to signaling within hHMAs, a dot plot of differential ion channel gene expression in hHMAs showed MPs express predominantly calcium and potassium ion channels, with high specificity for *CACNA1F*, *KCNA3*, *KCNJ10*, *KCNMA1*, *KCNN4*, *TRPA1*, *TRPC2*, *TRPM2*, *TRPV2*, and *TRPV4* (Figure 6H). Quantified values for action potential duration (APD) at 90%, 50%, and 30% repolarization (APD90, APD50, APD30) demonstrated significant differences between cardiomyocytes and MPs, with MPs displaying prolonged APD90, APD50, and APD30 as compared to cardiomyocytes (Figure 6I-L). Moreover, heatmap analysis of normalized log_2_FC pseudo-bulk gene expression data (Figure 6M) revealed upregulation of genes associated with Phase 2, 3, and 4 of the cardiac action potential in day 26 hHMAs when compared to hHOs. These findings suggest that MPs actively contribute to the electrophysiological activity and maturation of hHMAs.

### Chronic inflammatory conditions leading to prolonged NLRP3 activation increase the risk of atrial arrhythmias in hHMAs

Many clinical and animal studies have shown a connection between inflammation and arrhythmogenesis^76^, particularly the most common cardiac arrhythmia: atrial fibrillation. A critical missing component of this inflammatory response in prior *in vitro* models has been the lack of tissue-resident and recruited MPs. To demonstrate the pre-clinical value of our hHMA model, we sought to create a high-throughput *in vitro* inflammatory-hHMA platform to elucidate the cellular and molecular motifs that lead to atrial arrhythmia. We hypothesized that chronic activation of the NLRP3 inflammasome complex, which has been persistently associated with AF in humans^46^, could trigger hHMA remodeling and arrhythmicity. We designed a series of conditions (low, medium and high dose) involving the chronic administration of physiologically relevant^77,78^ concentrations of pro-inflammatory triggers (LPS, interferon-gamma, and interleukin-1β) common in chronic inflammatory conditions in human patients, and applied them to hHMAs. Day 20 hHMAs were matured using a combination of previously described methods (developmental maturation for 10 days, followed by cardiomyocyte maturation medium from day 30-64)^22^. hHMAs were exposed to pro-inflammatory factors from day 30 through day 64 of hHMA culture (Figure 7A). To screen for irregular rhythms, MUSCLEMOTION^79^ was used to generate contraction traces from phase contrast microscopy videos of Ctrl, Low, Med, and High exposure hHMAs (Figure S7A). Irregular rhythms were noted at day 64 and were analyzed with FluoVolt live-cell imaging. FluoVolt traces demonstrated that hHMAs exposed to pro-inflammatory cytokines had a higher beat rate than expected (tachycardia) when compared to Ctrl hHMAs (Figure 7B-C). Spontaneous irregular rhythms and cessation of FluoVolt signaling were quantified for each treatment group and those exposed to pro-inflammatory cytokines demonstrated significantly increased irregular rhythms as compared to Ctrl hHMAs (Figure 7D-E), which mirrored the MUSLCEMOTION contraction curves (Figure 7F). Moreover, quantification of maximum contraction amplitude across individual hHMAs under different conditions showed significant variability, with treated hHMAs exhibiting a reduced contraction amplitude (Figure 7E). hHMA morphological changes were also noted, with hHMAs in the High condition being significantly larger to Ctrl, Low, and Med conditions while remaining similar in circularity (Figure S7B-D; Videos S4-7). hHMAs in the Low, Med, and High conditions had noticeably more CD45+ expression than those in the Ctrl condition, suggesting tissue-resident macrophage expansion, a previously described phenomenon in atrial arrhythmias. Metabolically, although not statistically significant, hHMAs had a stepwise increase in oxygen consumption rate (OCR) with larger pro-inflammatory factor exposures (Figure S7E). Furthermore, action potential durations at 30% and 50% were significantly longer in High hHMAs versus Ctrl hHMAs (Figure 7H-J). Irregular electrophysiologic phenomena classically associated with atrial fibrillation, such as early afterdepolarizations (EADs), delayed after depolarizations (DADs), and spontaneous calcium elevations (SCaEs), were more prevalent in hHMAs exposed to pro-inflammatory factors (Figures 7K-M and S7F-K) than Ctrl hHMAs. The electrophysiological traces confirmed these events were predominantly present in atrial cardiomyocytes. To validate that this was due to NLRP3 activation as expected, RT-qPCR was performed on hHMAs in Ctrl, Low, Med, and High conditions at day 64 (Figure 7N). A dose-dependent, statistically significant increase in the expression of *NLRP3*, *CASP1*, *IL-1*β, *CD68*, *COL1A1*, *LGALS3*, and *RYR2* was detected, with larger pro-inflammatory factor exposures leading to higher activation in hHMAs. These proteins were significantly expressed in mammals’ hearts subject to chronic NLRP3 activation induced AF^46^. These results show that chronic activation of the NLRP3-inflammasome in hHMAs induces spontaneous arrhythmias and features consistent with AF and supports the pre-clinical value of our hHMA model for arrhythmia disease modeling and drug development in the near future.

**Figure 7.**
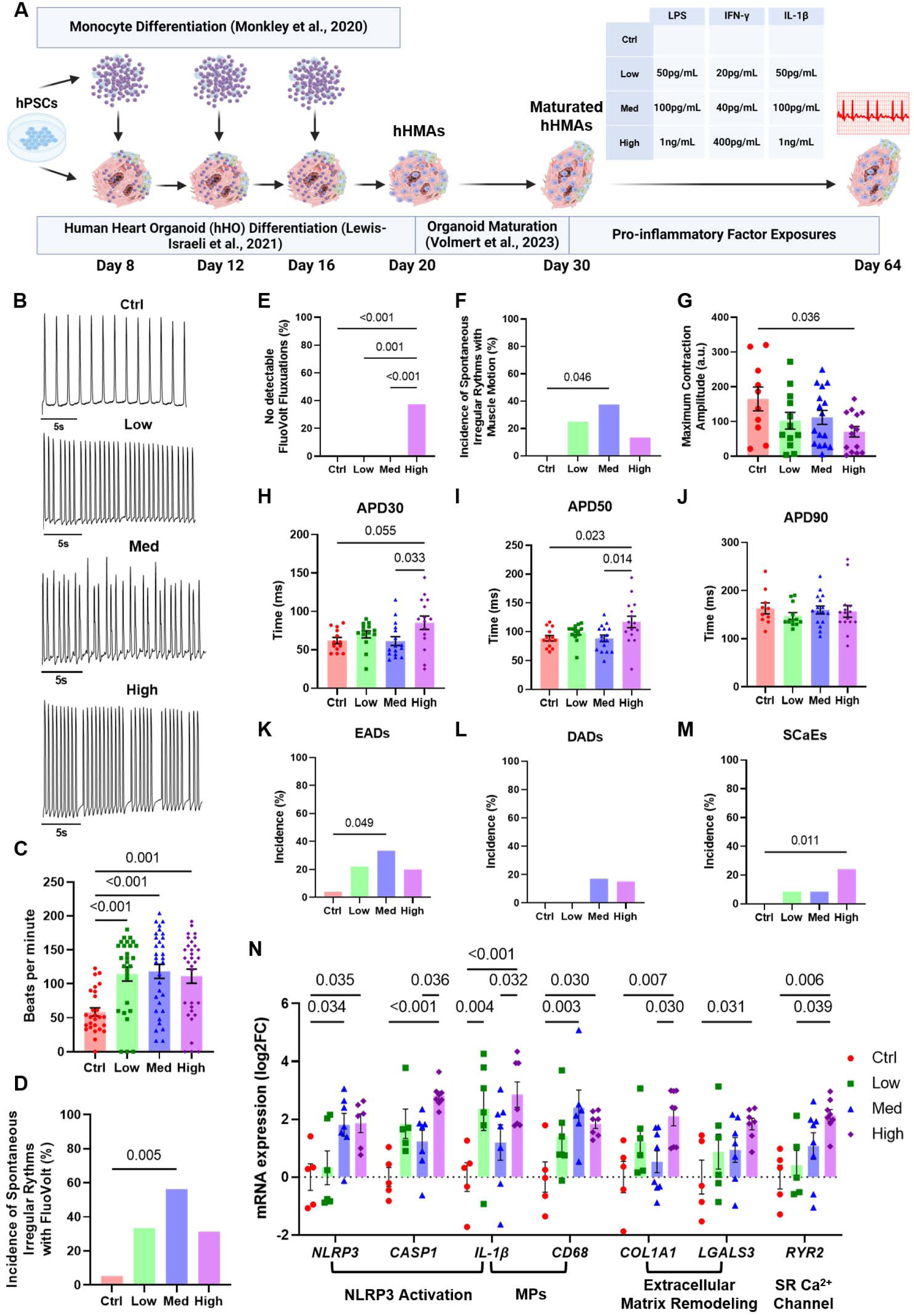
Prolonged NLRP3 activation results in arrhythmias in a proportion hHMAs with evidence of atrial fibrillation. (A) Schematic of the inflammatory-arrhythmia disease model in hHMAs. (B) Representative FluoVolt traces of Ctrl, Low, Med, and High hHMAs at high magnification. n ≥ 16. (C) Quantification of beat rate using the FIJI Muscle Motion on phase-contrast microscopy video recordings of day 64 Ctrl, Low, Med, and High hHMAs. n ≥ 27. (D) Quantification of the percentage of hHMAs with spontaneous irregular rhythms detected using FluoVolt live cell imaging of Ctrl, Low, Med, and High hHMAs. Irregular rhythms are defined as the presence of ectopic beats and irregular waveforms. n ≥ 16. (E) Quantification of undetectable FluoVolt fluctuations in Ctrl, Low, Med, and High hHMAs. n ≥ 15. (F) Quantification of the percentage of hHMAs with spontaneous irregular rhythms detected using the FIJI Plugin MuscleMotion live cell imaging of Ctrl, Low, Med, and High hHMAs. Irregular rhythms are defined as the presence of ectopic beats and inconsistent peak-to-peak distances. n ≥ 16. (G) Quantification of the maximum contraction amplitude for an individual hHMA in each condition. n ≥ 10. (H) Quantification of APD30s from action potentials in day 64 hHMAs from each condition using FluoVolt video recordings. n ≥ 13. (I) Quantification of APD50s from action potentials in day 64 hHMAs from each condition using FluoVolt video recordings. n ≥ 13. (J) Quantification of APD90s from action potentials in day 64 hHMAs from each condition using FluoVolt video recordings. n ≥ 13. (K) Quantification of the incidence of EADs in hHMAs amongst each condition. n ≥ 20. (L) Quantification of the incidence of DADs in hHMAs amongst each condition. n ≥ 20. (M) Quantification of the incidence of SCaEs in hHMAs amongst each condition. n ≥ 20. (N) RT-qPCR analyses on bulk RNA from day 64 hHMAs for proteins expressed in mammalian hearts with AF. Transcripts are representative in genes involved in the NLRP3 inflammasome activation pathway, MPs, extracellular matrix remodeling, and sarcoplasmic reticulum calcium handling. n ≥ 5. For graphs in (C-M): Value = mean± s.e.m., 1-way ANOVA multiple-comparison test. For graph in (N): Value = mean± s.e.m., 2-way ANOVA multiple-comparison test.

## DISCUSSION

In this study, we report the development of an integrated hHMA featuring autologous cardiac tissue-resident MPs based entirely on developmental self-organization. Our findings demonstrate that hHOs can be effectively populated with MPs through the integration of hPSC-derived monocytes, mimicking the developmental process of MP infiltration observed during embryonic heart development^58^. MPs in hHMAs not only highly resemble cardiac embryonic MPs at the transcriptome level when compared with human embryonic hearts^30,64^, but also persist over long periods of time in the hHMAs, which is consistent with the longevity of tissue-resident MPs in the heart^60^. Notably, the persistence and distribution of MPs in hHMAs align well with established *in vivo* data^30,64^, suggesting that this *in vitro* model recapitulates key aspects of early human heart development. Thus, our hHMAs provide a novel platform to isolate and study the effects of MPs on human cardiogenesis in a controlled setting.

The ability of tissue-resident MPs to influence cell communication within hHMAs was particularly striking. scRNA-seq data revealed that MP integration alters the composition of critical cardiac cell types, such as CFs, VCMs, and VCs. These are notable as MPs have been shown to modify heart valves in heart development and disease^80^, communicate extensively with CFs in the setting of cardiac wound healing and homeostasis^81^, and, in *in vitro* studies, preferentially induce a VCM phenotype in engineered heart tissues^41^. Our data suggested tissue-resident MPs acquire different roles in the heart and have association preferences to various cell types. At least three different subtypes could be identified, with a predilection for interstitial cells, cardiomyocytes, and chamber interior (Figure S5A-B). Further studies will be needed to identify specific sub-roles of these MP populations and how they impact the viability and cellular makeup of the developing heart.

It is well-established that tissue-resident MP signaling are essential for cell-to-cell communication in the heart in both homeostatic^82^ and disease^83,84^ contexts. We explored the transcriptome and EV proteome of hHMAs in profound detail to identify communication networks between MPs and other cardiac cell types. Our study provides a first-of-its-kind view of how MPs communicate with other cell types in a developmental human heart model. For example, LDHA induces the proliferation of cardiomyocytes in mice^85^, so it is notable that LDHA was explicitly expressed by VCMs and found exclusively in the EVs from hHMA (Figure 4G), suggesting a unique signaling pathway between MPs and VCMs that could induce VCM proliferation. Overall, these rich datasets will be crucial to uncovering cell-to-cell communication networks between MPs and the heart in the near future.

Regarding the role of MPs in cardiac conductance, our data suggests that MPs contribute to the electrophysiological properties of hHMAs by forming GAP junctions with cardiomyocytes and participating in calcium signaling and action potential propagation, confirming *in vivo* findings from past studies in rodents^28^. The formation of CX43-mediated GAP junctions between MPs and cardiomyocytes indicated functional coupling between these two cell types, allowing MPs to influence the electrical conductance of the heart. Live-cell imaging further confirmed that MPs exhibited synchronized calcium transients and action potentials with neighboring cardiomyocytes. Based on our observation that MPs in hHMAs exhibit more prolonged action potentials compared to cardiomyocytes (Figure 6I) and aligned with previous studies on MPs in cardiac electrophysiology^28^, we propose that MPs function similarly to capacitors, modulating action potential propagation, which is particularly important in the atrioventricular node. This effect would shorten action potential duration in MP-rich areas, increasing cardiomyocyte excitability. This would explain why a more significant MP presence, particularly in the AV-node, would lead to cardiac arrythmias like AF, as a recent study has shown^86^. Furthermore, with the upregulation of genes associated with calcium and potassium channels involved with cardiac action potentials (Figure 6H and 6M).

One of the most novel aspects of this study is the modeling of arrhythmogenesis in a human heart platform, with hHMAs, through the activation of the NLRP3 inflammasome. Prolonged NLRP3 activation in hHMAs resulted in increased arrhythmogenic events (Figure 7 and S7) such as EADs and DADs, SCaEs, irregular rhythms, and an elevated beat rate, all hallmarks of atrial fibrillation^42^. These data provide compelling evidence that MP-driven inflammation, mediated by the NLRP3 inflammasome, contributes to the pathogenesis of arrhythmias in human heart tissue. The molecular signatures observed in NLRP3-activated hHMAs, such as upregulation of genes involved in calcium handling, ECM remodeling, and MP activation, mirror those found in mammalian and human hearts with AF^46^. This suggests that hHMAs can be a powerful *in vitro* model for studying the underlying AF’s inflammatory mechanisms and testing potential therapeutic interventions targeting the NLRP3 inflammasome activation pathway. Future studies should utilize the hHMA model for drug discovery to discover optimized therapies for inflammation-induced arrhythmias in a human heart model. Based on the role MPs play in facilitating conductance in the heart and how inflammation induces cardiac arrhythmias^87^, it’s conceivable that immunotherapies may be an appropriate treatment option for AF in the presence of inflammation.

It is important to acknowledge that while hHMAs represent a significant advancement in modeling human heart development and disease, they remain an *in vitro* system that does not fully replicate the complexity of a fully developed human heart. Our hHMAs lack complete vascularization, physiologically relevant mechanical forces, and concrete cellular interactions present *in vivo* during heart development, such as with neural crest cells. These additional factors are critical for fully mimicking heart physiology. Additionally, although hHMAs provide valuable insights into the role of tissue-resident immune cell types in heart development, the absence of other systemic influences, such as other circulating immune cells and hormonal signals, limits the scope of some of our findings.

In conclusion, this study establishes hHMAs as a novel and physiologically relevant model for investigating the role of tissue-resident MPs in human heart development and disease. We demonstrated that MPs contribute to a wide range of processes in human cardiac tissue, including ECM remodeling, efferocytosis, cell-to-cell communication, and electrophysiology, underscoring their critical role in human heart homeostasis. Furthermore, the ability to model NLRP3-induced arrhythmogenesis in hHMAs provides new translational opportunities to explore the link between inflammation, MP activity, and arrhythmogenesis. This model holds promise for advancing our understanding of MP-driven cardiac pathologies and identifying new therapeutic strategies for cardiovascular diseases such as atrial fibrillation.

## METHODS

### Stem cell culture

The following human pluripotent stem cell lines were used for this study: ESC-H9, ESC-H9- mCherry, ESC-H1, hiPSC-L1. Pluripotency and genomic stability were tested for all hPSC lines used. hPSCs were cultured in Essential 8 Flex medium with 1% penicillin/streptomycin (Gibco) in 6-well plates on growth factor reduced Matrigel (Corning) inside an incubator at 37°C and 5% CO_2_. hPSCs were passaged using ReLeSR passaging reagent upon reaching 60-80% confluency. ESC-H9 derived hHOs and hHMAs were used for most studies in this report unless otherwise noted.

### Human monocyte differentiation

The monocytic lineage differentiation protocol is based on a 5 sequential step protocol with some minor adjustments^47,49^. The hPSC lines used were ESC-H9 or PSC-L1. To generate a more robust protocol with a higher yield of CD14+ cells, the protocol was modified by using 60 embryoid bodies (EBs) of known cell number (3x10^4^) per 6-well plate with 10 EBs per well instead of using 30 embryoid bodies (EBs) of known cell number (30x10^3^) per 6-well plate with 5 EBs per well. With these adjustments, at day 0, the differentiation and generation of EBs was started simultaneously by seeding undifferentiated single cell hPSCs at a concentration of 3x10^4^ cells per well in a 96 Ultralow attachment plate with round-bottom wells (Corning) in 100 μl media (E8 Flex complete with 2μM ROCK inhibitor (Thiazovivin) and 80 ng/ml BMP4. The plate was centrifuged at 200xg, for 3 min. at room temperature. On day 2, the EBs were transferred to growth factor-reduced Matrigel coated wells (Corning), five EBs per well in 6-well plates (Corning) in the same media described above. For the rest of the differentiation, the Yanagimachi protocol was followed^47^. Every four days, from day 18 to day 42, the suspension cells were positively sorted by MACS using CD14 + MicroBeads generating 7 batches of CD14+ monocytic lineage-directed cells. The Cao et al, 2019 protocol was adapted for MACS purification^48^. Yields from day 26 to day 42 were significantly larger than those from day 18 and day 22 collections. Thus, suspension cells between day 26 and day 42 were positively sorted by MACS using CD14 + MicroBeads and used for additions to human heart organoids.

### Self-assembling hHO differentiation

A detailed protocol which describes the generation and differentiation of human heart organoids is provided^17^. Human embryonic stem cell (hESC) line H9 (WiCell, WA09) was grown to 60-80% confluency on a 6-well plate and dissociated using Accutase (Innovative Cell Technologies) to obtain a single-cell solution. H9s were collected and centrifuged at 300 g for 5-minutes (min) and resuspended in Essential 8 Flex medium (Gibco) containing 2μM ROCK inhibitor (Thiazovivin) (Millipore Sigma). H9s were counted using a Moxi cell counter (Orflo Technologies) and 10,000 cells were seeded in 100µL per well in a round bottom 96 well ultra-low attachment plate (Costar). The plate was then centrifuged at 100 g for 3-min and subsequently placed inside a 37°C and 5% CO_2_ incubator (these same incubation conditions were used for the entire protocol). After 24 hours (h), 50 μL of medium was removed from each well and 200 μL of fresh Essential 8 Flex Medium with 1% penicillin/streptomycin was added to obtain a final volume of 250 μL per well. The plate was incubated for 24-h. For every media change onward, 166µL of medium was removed from each well and 166µL of the listed medium was added to each well. After incubation, medium was removed from each well and RPMI with B27 supplement without insulin (Gibco) supplemented with 1% penicillin/streptomycin (Gibco) (hereafter termed “RPMI/B27 minus insulin”) containing CHIR99021, BMP4, and Activin A was added to each well to obtain final concentrations of 4 μM CHIR99021, 36 pM (1.25 ng/mL) BMP4, and 8 pM (1.00 ng/mL) Activin A. The plate was subsequently incubated, and, after exactly 24-h, the medium was removed from each well and replaced with fresh RPMI/B27 minus insulin. 24-h later, spent medium was removed from each well and RPMI/B27 minus insulin with Wnt-C59 (Selleck) was added to obtain a final concentration of 2 μM Wnt-C59 inside each well. The plate was then incubated for 48-h. Afterwards, the medium was removed and replaced with fresh RPMI/B27 minus insulin and incubated for another 48-h. Following the incubation, spent medium was removed and replaced with RPMI with B27 supplement (with insulin) and 1% penicillin streptomycin (hereafter termed RPMI/B27). The plate was incubated for 24-h. Afterwards, medium was removed from each well and RPMI/B27 containing CHIR99021 was added to obtain a final concentration of 2 μM CHIR99021 per well. The plate was incubated for 1 hour. After 1-h, the medium was removed from each well and fresh RPMI/B27 was added to each well. The plate was incubated for 48-h. From days 9 to 20, every 48-h, media changes were performed by removing media from each well and adding fresh RPMI/B27.

### Generation of hHMAs

On day 8 of hHO differentiation, CD14-MACS purified monocytes were added to human heart organoids. In more detail, before collecting the cells from Day 26 through Day 42 of monocyte factory culture. Prewarm FACS buffer (0.5% BSA and 2mM EDTA in PBS, pH=7.4) to room temperature. Prepare enough Step 4 Medium^47^ so that 3mL of Step 4 Medium can replenish each monocyte factory well. To dislodge any unattached monocytes, pipet spent monocyte factory medium very gently in each well up and down no more than two times using a 1-ml pipet and collect the whole cell suspension in a 50-ml tube. Make sure to not pipet so hard that the monocyte factories become damaged. Quickly, add Step 4 Medium to the monocyte factories. After collecting the monocytes, wash off any lingering CD14 Microbeads from the monocytes and equilibrate the monocytes to organoid medium, gently resuspend pelleted monocytes in 10mL of RPMI +ins +B27 medium, centrifuge at 300xg for 3 min at room temperature and discard the supernatant. Repeat this step two more times. After the last centrifuge spin, resuspend CD14+ cells in 1ml of RPMI +ins +B27 +P/S medium and calculate the volume of monocytes needed to add 20,000 monocytes per organoid using RPMI +ins +B27 as the diluent. On day 10, day 14, and day 18, perform an RPMI +ins +B27 media change. On days 12 and 16, repeat the monocyte addition. By day 20, MP integrated organoids were ready for analysis.

### Organoid and assembloid dissociation

Culture medium was carefully removed and discarded from the hHO. The hHO was washed once with PBS to remove any loosely attached cells. RPMI+B27 with 2% BSA (w/v) was prepared and filtered to make sterile. Cardiomyocyte Dissociation Medium was warmed to 37°C prior to dissociation. 500µL of the Cardiomyocyte Dissociation Medium was applied to the organoid and shaken at 37°C at 250rpm for 5-min. The dissociation medium was then collected from the organoid containing RPMI+B27 with 2% BSA. The dissociation medium addition, incubation, and collection was repeated until 20- min passed. The collection was filtered with a 40µm filter and then centrifuged at 300g for 5min. The supernatant was decanted, and the pelleted cells were resuspended in a smaller volume of FACS buffer for flow cytometry or RPMI+B27 + 2% BSA for scRNA-seq. After the cells were resuspended, the cells were filtered again with a 40µm filter.

### Flow cytometry

Following CD14 MACS purification, monocytes were washed once with FACS Buffer and blocked with Human BD Fc Block^TM^ (BD Biosciences) for 10-mins at room temperature. Cells were then stained with CD45-FITC (HI30; BD Biosciences) and CD14-AF647 (63D3.rMAb; BD Biosciences) or CD163 (MOPC-21; BD Biosciences) at 4°C for 30-min in the dark. Stained single-cell suspensions were washed twice with FACS buffer, resuspended in 300µL, and filtered through a 100µm cell strainer. Cell debris was excluded from the analysis with gating from the FSC-SSC plot and singlets were selected from FSC-A vs FSC-H and SSC-A and SSC-H plots. For dissociated organoids, the same protocol was followed except singlets were not ruled out during the analysis. Live/cell staining was accomplished with 7-amino-actinomycin a (7-AAD), demonstrating >90% viability in all dissociated organoid samples. MPs were gated for CD45-FITC cells vs SSC-A as inspired from previous work^58^.

### Immunofluorescence staining

The immunofluorescence staining protocol has been described previously^17,22^. hHOs were transferred from the round bottom ultra-low attachment 96 well plate to 1.5 mL microcentrifuge tubes (Eppendorf) using a cut 200 μL pipette tip (to increase tip bore diameter as to not disturb the organoid). Organoids were washed one time with PBS to remove any loosely attached cells. Organoids were fixed in 4% paraformaldehyde (VWR) in PBS for 30-min. Following, organoids were washed using PBS- Glycine (1.5 g/L) three times for 5-min each. Organoids were then blocked and permeabilized using a solution containing 10% Donkey Normal Serum (Sigma), 0.5% Triton X- 100 (Sigma), and 0.5% BSA (Thermo Fisher Scientific) in PBS on a thermal mixer at 300rpm at 4 °C overnight. Organoids were then washed 3 times using PBS and incubated with primary antibodies (Supplementary Table 1) within a solution containing 1% Donkey Normal Serum, 0.5% Triton X-100, and 0.5% BSA in PBS (hereafter termed “Antibody Solution”) on a thermal mixer at 300rpm at 4 °C for 24-h. Following, organoids were washed 3 times for 5-min each using PBS. Organoids were then incubated with secondary antibodies (Supplementary Table 1) in Antibody Solution on a thermal mixer at 300rpm at 4°C for 24-h in the dark. Subsequently, organoids were washed 3 times for 5-min each using PBS and mounted on glass microscope slides (Fisher Scientific). 90 μm Polybead Microspheres (Polyscience, Inc.) were placed between the slide and a No. 1.5 coverslip (VWR) to provide support pillars such that the organoids could retain three dimensionality. Organoids were transferred to the glass microscope slides using a cut 200µL pipette tip and mounted using a clearing solution described previously^88^.

### Confocal microscopy and image analyses

Immunofluorescence images were acquired using a Nikon Instruments A1 Confocal Laser Microscope. Images were analyzed using Fiji. For cell quantification of TNNT2, MYL3, and CD45 area was determined by quantifying the total positive pixel signal for each antigen and dividing by the total pixel shadow of the organoid for a given z-stack. This was done for a minimum of 3 z-stack images per organoid and averaged amongst each other to generate a percentage of area that is positive for a given protein. The I/E ratio (Figures 1D and S1G) was determined by quantifying CD45+ area in z-stacks inside of the organoids (interior) and dividing the CD45+ area in in z-stacks that are on the surface (exterior) of the organoids. Sarcomere length was determined by measuring the distance between z-lines in high-magnification immunofluorescence images using FIJI (Figure 5E). The region of interest (ROI) tool was used to trace the CALR+ areas in hHMAs (Figure 5J). CD45+ signal that was within the CALR+ region of interest was considered (CALR+ area) and CD45+ signal that was outside the CALR ROI was considered (CALR- area). MERTK puncta quantifications were done by counting the number MERTK+ puncta within CD45+ cells as compared to other cells in frame of the high-magnification image (Figure 5K). Nuclei (DAPI+ areas) were quantified and assigned to CD45+ Cells or Other Cells for the analyses. Furthermore, the Fiji ROI tool was used to outline TNNT2+, WT1+, or hollow regions (cavities) within a minimum of 3 z-stacks per organoid (Figure S5B). CD45+ signal was categorized into TNNT2+ area, WT1+ area, or cavities if it fell within these respective ROIs. The CD45+ signal was then summed and used as the denominator for each categorization, providing a CD45+ Cells % for each ROI.

### qRT-PCR

Following CD14-MACS purification, monocytes were collected, pelleted, and preserved in RNAprotect (Qiagen) at −20°C. Organoids were harvested on day 20 and similarly stored in RNAprotect at - 20°C. RNA extraction was conducted using the Qiagen RNEasy Mini Kit, mostly following the manufacturer’s protocol. Organoid samples were lysed with the Bead Mill 4 Homogenizer (Fisher Scientific) at speed setting 2 for 30 seconds. RNA concentration was determined using a NanoDrop One (Thermo Fisher Scientific), and only samples with a concentration of at least 10 ng/μL proceeded to reverse transcription. cDNA synthesis was performed using the Quantitect Reverse Transcription Kit (Qiagen) and stored at −20°C. Primers for real-time qPCR were designed with the Primer Quest tool (Integrated DNA Technologies). SYBR Green (Thermo Fisher Scientific) served as the DNA-intercalating dye for the reaction vessel. Real-time qPCR was conducted on the QuantStudio 5 Real-Time PCR system (Applied Biosystems) with a total reaction volume of 20 μL. Gene expression was normalized to HPRT1 for each sample, and fold change was calculated using the double delta CT method. mRNA expression data are presented as fold change (FC) and/or log2FC relative to the Control.

### scRNA-seq

Libraries were prepared using the 10x Chromium Next GEM Single Cell 3’ Kit, v3.1 and associated components. Completed libraries were QC’d and quantified using a combination of Agilent 4200 TapeStation HS DNA1000 and Invitrogen Collibri Library Quantification qPCR assays. Libraries from each pool were pooled in equimolar proportions, and the pool was quantified again using the Invitrogen Collibri qPCR assay. These two project pools were then pooled in equal proportions and this final pool again quantified by qPCR. The pool was loaded onto 1 lane of an Illumina NovaSeq 6000 S4 flow cell (v1.5) and sequencing was performed in a 2x150bp paired end format using 300 NovaSeq v1.5 reagent cartridge. Base calling was done by Illumina Real Time Analysis (RTA) v3.4.4 and output of RTA was demultiplexed and converted to FastQ format with Illumina Bcl2fastq v2.20.0. For Bcl2fastq only the first 28 bases of read #1 and 100 bases of read #2 were output. The first 28bp of read 1 includes the 10x cell barcodes and UMIs, read 2 is the cDNA read. Demultiplexed fastq files, were processed using 10x Genomics Cellranger count (v7.1.1) software using standard protocol including introns and exons data using Ensemble Homo_sapiens.GRCh38.110 genome. 10x count matrices were processed using standard Seurat (v5.0.2)^89^ pipeline and scCustomize^91^ helper functions. Cells were slightly filtered (nFeature_RNA > 2000, nFeature_RNA < 7500, nCount_RNA > 1500, nCount_RNA < 40000, percent_mito < 15). Samples were separately estimated to Cell Cycling and scaled by ’S.Score’ and ’G2M.Score’. To avoid over-integration and keep heterogeneity samples were integrated using fast integration using reciprocal PCA (RPCA) by Satijalab guidelines. UMAP dimensional reduction was generated with Louvain algorithm using the built-in Seurat implementation. Integration with published data was performed using Harmony (v0.1 Broad)^92^ on SCT transformed data, with individual dataset counts regressed using (percent_mito, percent_ribo, S.Score, G2M.Score, nCount_RNA, nFeature_RNA). Enrichr^90^ and Hallmark^93,94^ were used to assess gene ontologies. When Feature plots are presented, the color intensity represents the relative gene expression value per gene. Liana^95^ was used to elucidate ligand-receptor relationships and scDiffCom^96^ was used to identify differential ligand-receptor relationships between hHMAs and hHOs. Phantasus^97^ was used for hierarchal clustering and pseudobulk heatmap analyses.

### Live-cell imaging for calcium and voltage recordings

Intracellular Ca^2+^ and ion signaling was visualized using Fluo4 and FluoVolt dyes, respectively, imaged and analyzed as previously described^22^. Both dyes were prepared via the manufacturer’s instructions. Additionally, NucBlue was used to visualize cell nuclei. NucBlue was prepared by adding two drops per milliliter to basal media (RPMI +ins). hHOs were washed twice using 166 μL of RPMI +ins, then 166 μL of the NucBlue solution was added to achieve a final concentration of 100 nM. Organoids were incubated for 30 min at 37 °C and 5% CO_2_. Organoids were then washed twice using basal medium and transferred to a chambered coverglass slide (Cellvis) using a truncated 200 μL pipette tip. Images were acquired using a Cellvivo microscope (Olympus IX83) under normoxic culture conditions at 37°C. Samples were excited with 360 nm (for NucBlue), 488nm (for Fluo4/FluoVolt), and 594nm (for mCherry-MPs) light. Live-cell imaging videos were collected for at least 20 seconds at a frame rate of 100 frames/second. Data was processed using the Fiji and analyzed using the multi-measure tool, as described^22^. Samples were excited at 494 nm excitation and emissions were collected at 506nm. Data was processed using Fiji. Baseline F_0_ of fluorescence intensity was calculated using the lowest 50 intensity values in the acquired dataset. Fluorescence change ΔF/F_0_ was calculated using the equation:

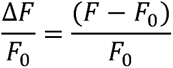

EADs, DADs, and SCaEs were quantified by identifying the presence of at least one of these phenomena in one hHMA video file.

### EV collection and isolation

Culture medium was collected from day 26 hHOs and hHMAs across one 96-well plate, totaling approximately 15 mL of medium containing EVs per sample. The medium was centrifuged at 400 x g for 10-min to remove large debris, then at 2,000 x g for 30-min to eliminate large particles, such as apoptotic bodies. To isolate the EVs, the media was placed into ultracentrifuge tubes, balanced with PBS, and centrifuged at 100,000g for 90-min at 4°C. After removing the supernatant, the pellet was resuspended in 1 mL of PBS. The sample volume was adjusted to two-thirds of the ultracentrifuge tube, and the sample was spun again at 100,000g for 90-min at 4°C. Finally, the supernatant was discarded, and the pellet was resuspended in 1 mL of PBS.

### Optical coherence tomography

To perform OCT imaging, we used a Spectral-Domain Optical Coherence Tomography (SD-OCT) system that is similar to those in our previous studies^22,98,99^. We used a superluminescent diode (EXALOS, EXC250023-00) as the light source, with a center wavelength of ∼1300 nm and a 3 dB spectrum range of ∼180 nm. The spectrometer (Wasatch Photonics, Cobra 1300) featured a 2048- pixel InGaAs line-scan camera (Sensors Unlimited, GL2048) and achieved a maximum A-scan rate of 147 kHz. Imaging involved a 5X objective lens with transverse and axial resolutions of ∼2.83 μm and ∼3.04 μm, respectively. We imaged and analyzed sixteen fixed Day 26 organoids from each group. We used customized MATLAB code to re-scale OCT images for isotropic pixel sizes in the x, y, and z dimensions. Amira software (Thermo Fisher Scientific) and ImageJ was employed for segmentation, 3D rendering and other visualization. Total volume and internal cavities of the organoids were quantified from the manual segmentation using Amira.

### LC-MS proteomics

For proteolytic digestion, SDS was added to the EV samples to 4% (w/v) and samples were digested overnight using S-traps (www.protifi.com) according to manufacturers’ instructions, using trypsin added to 500ng. After digestion, peptides were eluted from the S-trap, dried in a vacuum centrifuge and frozen at −20°C. Digested samples were re-suspended in 2% acetonitrile (ACN)/0.1% trifluoroacetic acid (TFA) to 20µL. An injection of 10µL was automatically made using a Thermo (www.thermo.com) EASYnLC 1000 onto a Thermo Acclaim PepMap RSLC 0.1mm x 20mm C18 trapping column and washed for ∼5min with buffer A. Bound peptides were then eluted over 35min onto a Thermo Acclaim PepMap RSLC 0.075mm x 150mm resolving column with a gradient of 5%B to 19%B from 0min to 19min, 19%B to 40%B from 19min to 24min and 40%B to 90B% from 24min to 25min. After the gradient, the column was washed with 90%B for 10min (Buffer A = 99.9% Water/0.1% Formic Acid, Buffer B = 80% Acetonitrile/0.1% Formic Acid/19.9% Water) at a constant flow rate of 300nl/min. Eluted peptides were sprayed into a Thermo Scientific Q-Exactive mass spectrometer (www.thermo.com) using a FlexSpray spray ion source. Survey scans were taken in the Orbi trap and the top ten ions in each survey scan are then subjected to automatic higher energy collision induced dissociation (HCD). The resulting MS/MS spectra are converted to peak lists using Mascot Distiller, v2.8.5 (www.matrixscience.com) and searched against a reference protein database containing all human sequences available from Uniprot (www.uniprot.org, downloaded 04-18-2023) appended with common laboratory contaminants (downloaded from www.thegpm.org, cRAP project) using the Mascot2 searching algorithm, v 2.8.3. The Mascot output was then analyzed using Scaffold, v5.3.3 (www.proteomesoftware.com) to validate protein identifications probabilistically. Assignments validated using the Scaffold 1% FDR confidence filter are considered valid. Proteomics results for each group were entered into the STRING database to create a STRING network utilizing evidence and medium confidence of 0.4. The desired biological and molecular processes determined on the protein network from gene ontology were presented. Mascot parameters for all databases were as follows: allow up to 2 missed tryptic sites, fixed modification of carbamidomethyl cysteine, variable modification of oxidation of methionine, peptide tolerance of +/- 10ppm, MS/MS tolerance of 0.02 Da, and FDR calculated using randomized database search.

### Generation of arrhythmogenic hHMAs with pro-inflammatory factors

Day 20 hHMAs were subjected to EMM2/1 maturation conditions up until day 30 as described before^22^. Day 30 hHMAs were then switched to MM maturation media^22^ and cultured in the presence of low (Low: 50pg/mL LPS, 20pg/mL IFN-γ, and 50pg/mL IL-1β), medium (Med: 100pg/mL LPS, 40pg/mL IFN-γ, and 100pg/mL IL-1β), or high (High: 1ng/mL LPS, 400pg/mL IFN-γ, and 1ng/mL IL-1β) concentrations of proinflammatory factors until Day 64, exchanging 166µL of media every 2-3 days.

### Phase contrast-microscopy and contraction analyses

Phase contrast-microscopy images were captured on an inverted Olympus bench-top light microscope and an inverted Leica Thunder Microscope. For live video of arrhythmogenic hHMAs, 1-minute-long videos were captured at a frame rate of at least 15 frames per second and maintained in normoxic conditions at 37°C on the Leica Thunder Microscope. The beat rate was determined by counting the total beats within the 1-minute-long video file. The FIJI plugin MUSCLEMOTION^79^ was used to determine contraction amplitude. The generated maximum contraction amplitude for each hHMA phase-contrast video was plotted and used in statistical comparisons.

## Statistics and reproducibility

All analyses were performed using GraphPad 9 software. All data presented a normal distribution. Statistical significance was evaluated with a standard unpaired Student t-test (2-tailed; P < 0.05) when appropriate. For multiple-comparison analysis, 1-way ANOVA was applied when appropriate (P < 0.05). The specific number of independent organoids used and the number of independent experiments (batches) for every experiment are indicated in figure legends. If not explicitly stated, images and data presented were from organoids derived from the hESC H9 cell line.

## Data availability

Bulk RNA-seq and scRNA-seq data sets have been deposited in the National Center for Biotechnology Information Gene Expression Omnibus repository under accession numbers GSE153185 and GSE280807, respectively. Proteomics data sets have been deposited in the Metabolomics Workbench repository under accession MSV000096319. All data generated or analyzed in this study are provided in the published article and its supplementary information files or can be obtained from the corresponding author upon request.

## Supporting information

Key resources table

Supplementary video descriptions

## Acknowledgments

We thank the IQ and MSU Advanced Microscopy Cores for access to confocal microscopes and the MSU Genomics Core, Stem Cell Core, and Proteomics Core for sequencing, cell culture, and mass spectrometry services, respectively. We also want to thank all members of the Aguirre Lab for their valuable comments and advice. Work in Dr. Aguirre’s laboratory was supported by startup funds from MSU, the National Institutes of Health (NIH) under award numbers K01HL135464, R01HL151505, by the American Heart Association under award numbers 19IPLOI34660342, 23IPA1053441, by the Corewell-MSU Foundation, Corewell Health, and the Alternatives Research and Development Foundation (ARDF). Work in Dr. Park’s lab was supported by startup funds from MSU and by the NIH under award number R01AR083086. Work in Dr. Contag’s lab was supported by the NSF under award number DE1001001. Work in Dr. Ashammakhi’s lab was supported by the American Heart Association under award number 23IPA1053441. Research in Dr. Zhou’s laboratory was supported by NIH grants R01-EB025209, R01-HL156265, and R21EB03268401A1.

## Author Contributions

C.O. and A.A. designed all experiments and conceptualized all the work. C.O. and A.A. assembled the figures and wrote the manuscript. C.O., S.C., M-L.S., M.D., P.M., H.B., M.S., A.H., and G.B. performed cell, organoid, and monocyte differentiations/culture. C.O. and M.P. performed flow cytometry experiments. C.O. and B.V. performed live-cell imaging experiments. S.A., I.N-R., N.A. and C.C. performed, analyzed and conceptualized proteomic and EV experiments. A.K. and S.P. integrated and processed raw scRNA-seq data. C.O. made cluster assignments and performed scRNA-seq analyses. N.C. helped conceptualize the arrhythmogenic hHMA experiments and model system. F.W. and C.Z. performed optical coherence tomography experiments and data analyses. W.C. and C.O. performed image analyses. C.O. performed all other experiments not explicitly mentioned in this section.

