## Supplementary figures for "Human heart assembloids with autologous tissue-resident macrophages recreate physiological immuno-cardiac interactions"

**Lead contact**

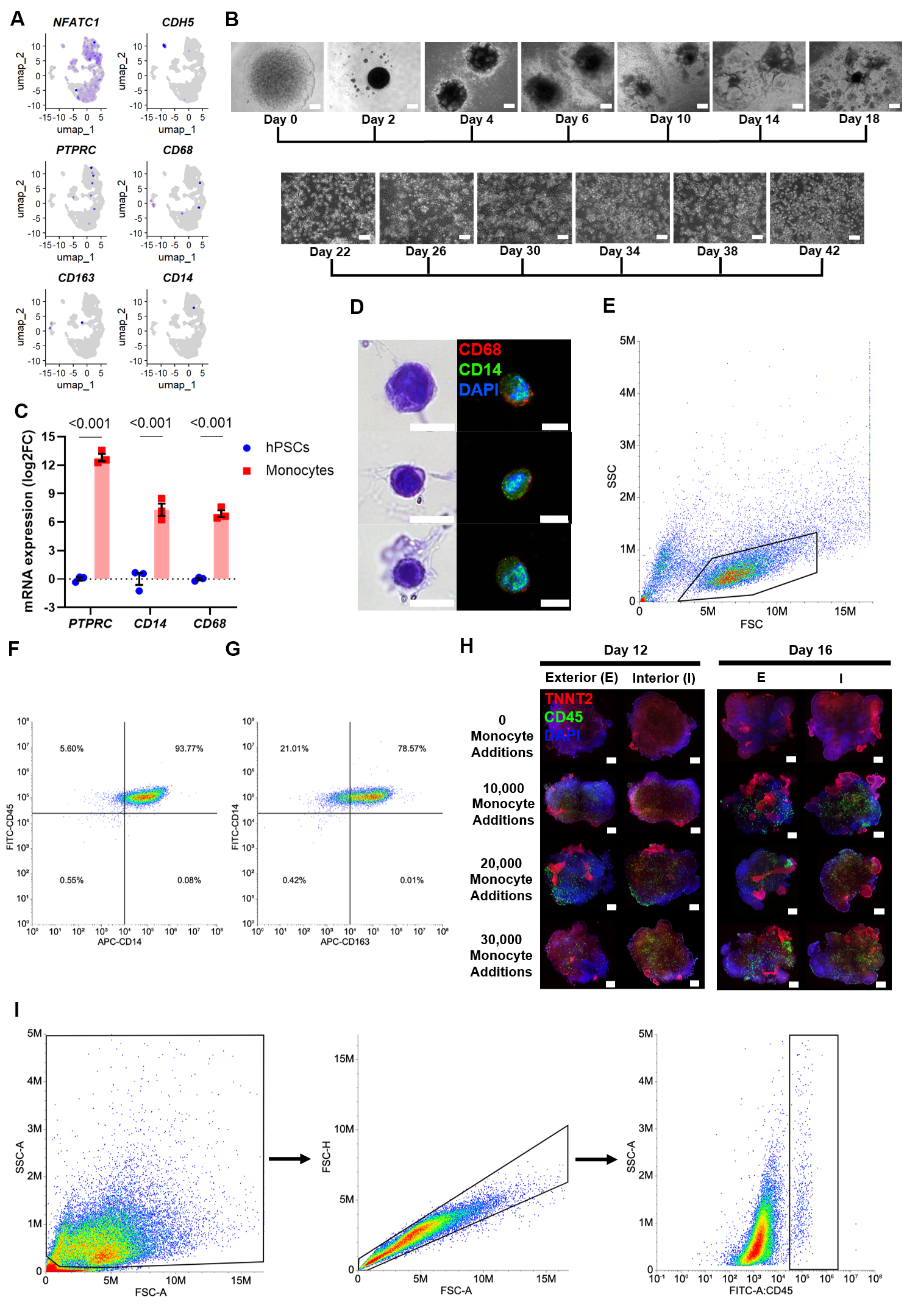

**Supplementary Figure 1. Differentiation of CD14+ embryonic monocytes from hPSCs and generation of hHMAs.**

(A) UMAP feature plots displaying relative expression for cardiac embryonic tissue-resident macrophages (MPs) markers, NFATC1, CDH5, PTPRC (CD45), CD68, CD163, and CD14 in hHOs (day 15 and day 26 hHO overlay). Color intensity represents the relative value of gene expression per gene.

(B) Timeline of phase-contrast microscopy images of monocyte factories over 42 days. Top row scale = 100µm. Bottom row scale = 50µm.

(C) RT-qPCR analyses on bulk RNA from PSCs and MACS-isolated CD14+ monocytes for monocyte/macrophage receptors, CD45, CD14, and CD68. n = 3. Value = mean ± s.e.m., Two-way ANOVA with multiple comparison test.

(D) Representative H&E staining (left) and immunofluorescence confocal microscopy images for CD68 (red), CD14 (green), and DAPI (blue) of MACS-isolated CD14+ monocytes. n = 3.

(E) FSC-SSC plot of CD14+ MACS purified cells that were added to hHOs. Black gate represents the cells analyzed in Figures S1F-G.

(F) Representative flow cytometry plots of CD14+ MACS purified cells gated for CD45-FITC and CD14-AF647. n = 3.

(G) Representative flow cytometry plots of CD14+ MACS purified cells gated for CD14-AF647 (left) or CD163-AF647 (right) staining. n = 3.

(H) Representative confocal IF images displaying the exterior (left panel) and interior (right panel) of day 12 and day 16 hHMAs stained for DAPI (blue), CD45 (green) and TNNT2 (red). Scale = 200µm. n ≥ 6.

(I) Gating strategy for identification of immune cells (CD45+) in dissociated hHMAs.

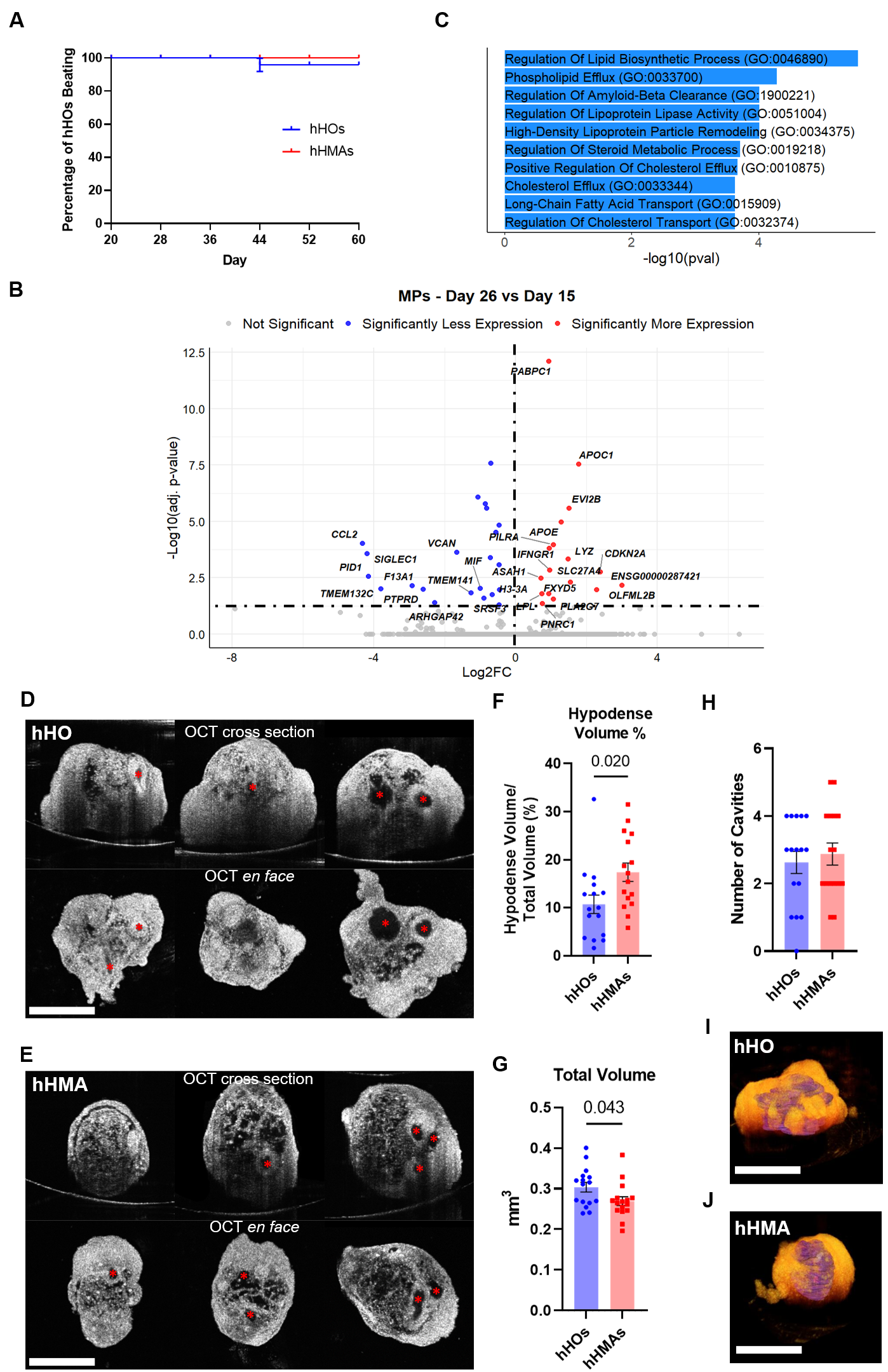

**Supplementary Figure 2. hHOs exhibit no endogenously derived MP-like populations and MPs alter the interior structure of hHMAs.**

(A) Quantification of control hHOs (blue) and hHMAs (red) visibly beating under live phase-contrast microscopy at day 20, 28, 36, 44, 52, and 60. n = 8. For all graphs: Value = mean ± s.e.m.

(B) Volcano plot displaying differential gene expression between Day 26 hHMAs and Day 15 hHMAs. Labels are present for significantly differentially expressed genes that are not mitochondrial or ribosomal. Transcripts that are significantly more abundant are red on the plot and those that are significantly less abundant are blue.

(C) Top 10 Gene ontology processes of 50 differentially expressed genes in MPs in Day 26 hHMAs versus Day 15 hHMAs.

(D – E) Representative OCT images of day 26 hHOs (D) and hHMAs (E). Top panels demonstrate the cross-sectional views while the bottom images represent en face view (top view). Red asterisks represent quantified isolated cavities. n = 16. Scale = 500µm.

(F) Quantification of hypodense OCT regions within the total volume of day 26 hHOs and hHMAs, represented as a percentage. n = 16.

(G) Quantification of the total volume of day 26 hHOs and hHMAs using OCT. n = 16.

(H) Quantification of the number of cavities within day 26 hHOs and hHMAs using OCT. n = 16.

(I) Representative 3D rendering of a day 26 hHO using OCT. n=16. Scale = 500µm.

(J) Representative 3D rendering of a day 26 hHMA using OCT. n=16. Scale = 500µm.

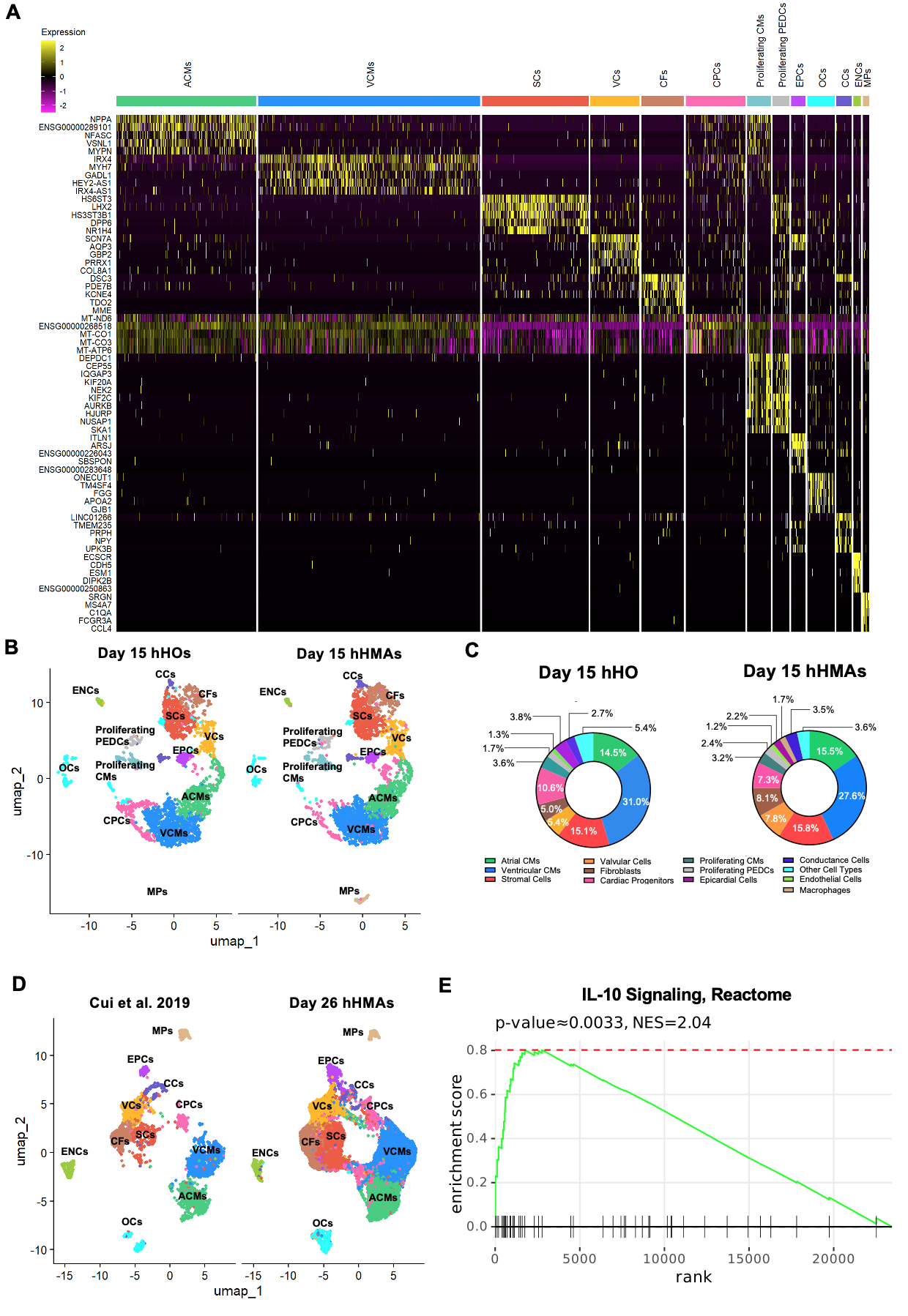

**Supplementary Figure 3. hHMA single cell transcriptome reveals similarities to embryonic human hearts.**

(A) Differential expression heatmap displaying the top 5 differentially expressed genes for all clusters.

(B) UMAP dimensional reduction plots of integrated scRNAseq data for each condition (from left to right): Day 15 hHOs and Day 15 hHMAs. Cluster identity labels are on the UMAP plots.

(C) Quantification of total cell count percentages per cluster for Day 15 hHOs and Day 15 hHMAs. Colors of regions correspond to the adjacent legend.

(D) UMAP dimensional reduction plots of integrated scRNAseq data from embryonic human hearts-Cui et al. 2019 (left) and Day 26 hHMAs (right). Cluster labels from Day 26 hHMAs are preserved from Fig 2A (right). Cui et al. 2019 UMAP displays overlayed cluster labels from Day 26 hHMAs (left).

(E) Gene set enrichment plot of differential gene expression between hHMAs and hHOs for IL-10 Signaling, Reactome.

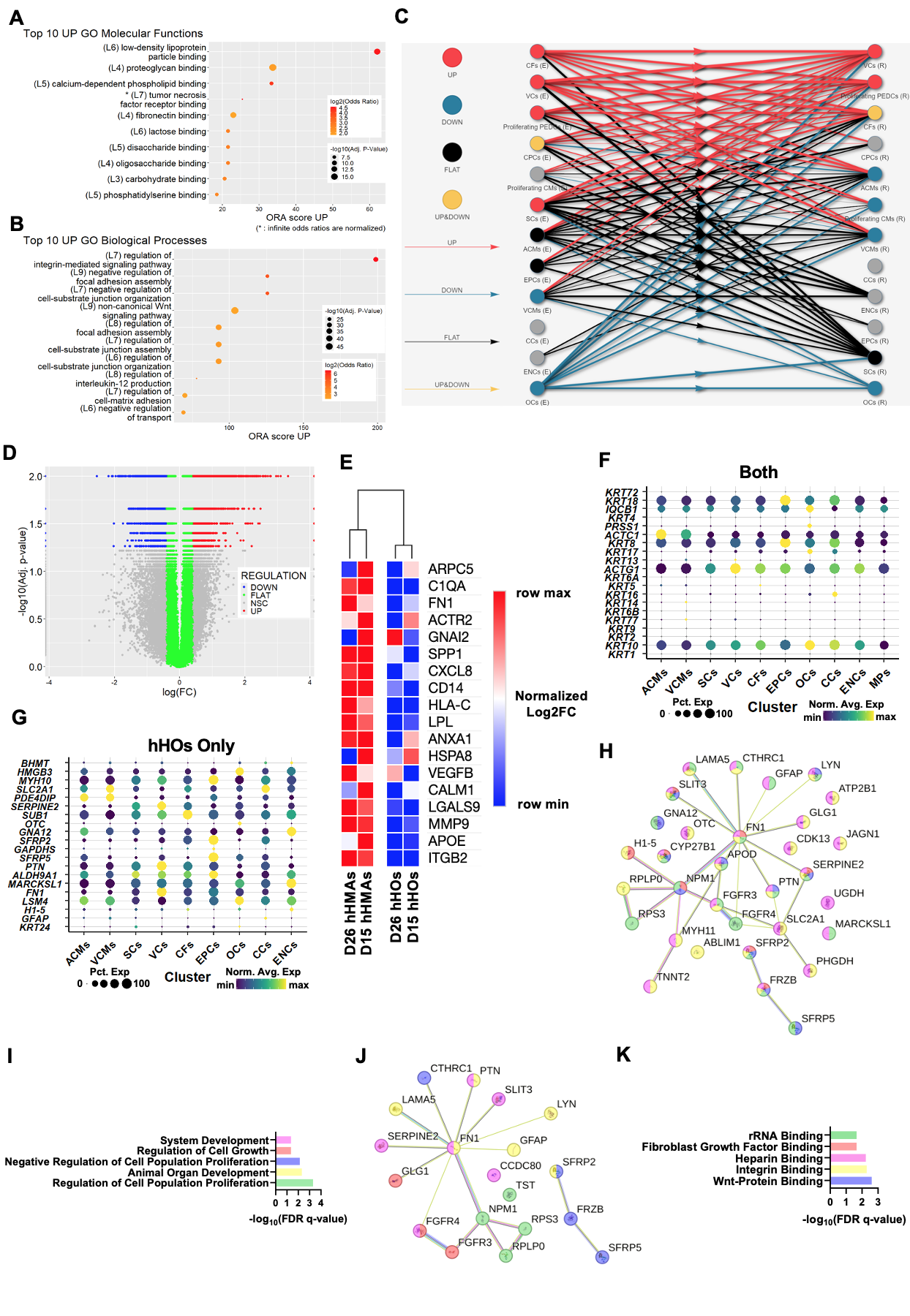

**Supplementary Figure 4. Cardiac tissue-resident MPs contribute critically to cell-cell signaling in human heart organoids.**

(A) Top 10 upregulated gene ontology molecular functions from the differential ligand-receptor analysis between day 26 hHMAs vs day 26 hHOs.
(B) Top 10 upregulated gene ontology biological processes from the differential ligand-receptor analysis between day 26 hHMAs vs day 26 hHOs.
(C) Day 26 hHMA vs hHO cell-type differential ligand-receptor analysis plot displaying upregulated (UP, red), downregulated (DOWN, blue), unchanged (Flat, black), and both up/downregulated (UP/DOWN, yellow) aggregated interactions from effector cell-types (E) to receiving cell-types (R).
(D) Volcano plot representing differential ligand-receptor interactions that are significantly upregulated (UP), downregulated (DOWN), unchanged (FLAT), and not significantly changed (NSC) due to the presence of MPs in day 26 hHMAs versus day 26 hHOs.
(E) Heatmap of normalized log2FC displaying expression of the most represented MP ligands across D15 hHOs, D15 hHMAs, D26 hHOs, and D26 hHMAs.
(F) Dot plot displaying differential expression of the 20 most prevalent proteins in EVs from both hHOs and hHMAs by cluster. Color is indicative of the normalized average gene expression between each cluster and the size of the circle is indicative of the percentage of cells within the cluster that express the respective gene.
(G) Dot plot displaying differential expression of the 20 most prevalent proteins in EVs from only hHOs by cluster. Color is indicative of the normalized average gene expression between each cluster and the size of the circle is indicative of the percentage of cells within the cluster that express the respective gene.
(H) String plot of proteins found in hHO EVs involved in the GO Biological Processes in I. Colors correlate to the color of bars found in I.
(I) 10 significantly upregulated GO Biological Processes using the proteins found in hHO EVs.
(J) String plot of proteins found in hHO EVs involved in the GO Molecular Functions involved in K. Colors correlate to the color of bars found in K.
(K) 10 significantly upregulated GO Molecular Functions using the proteins found in hHO EVs.

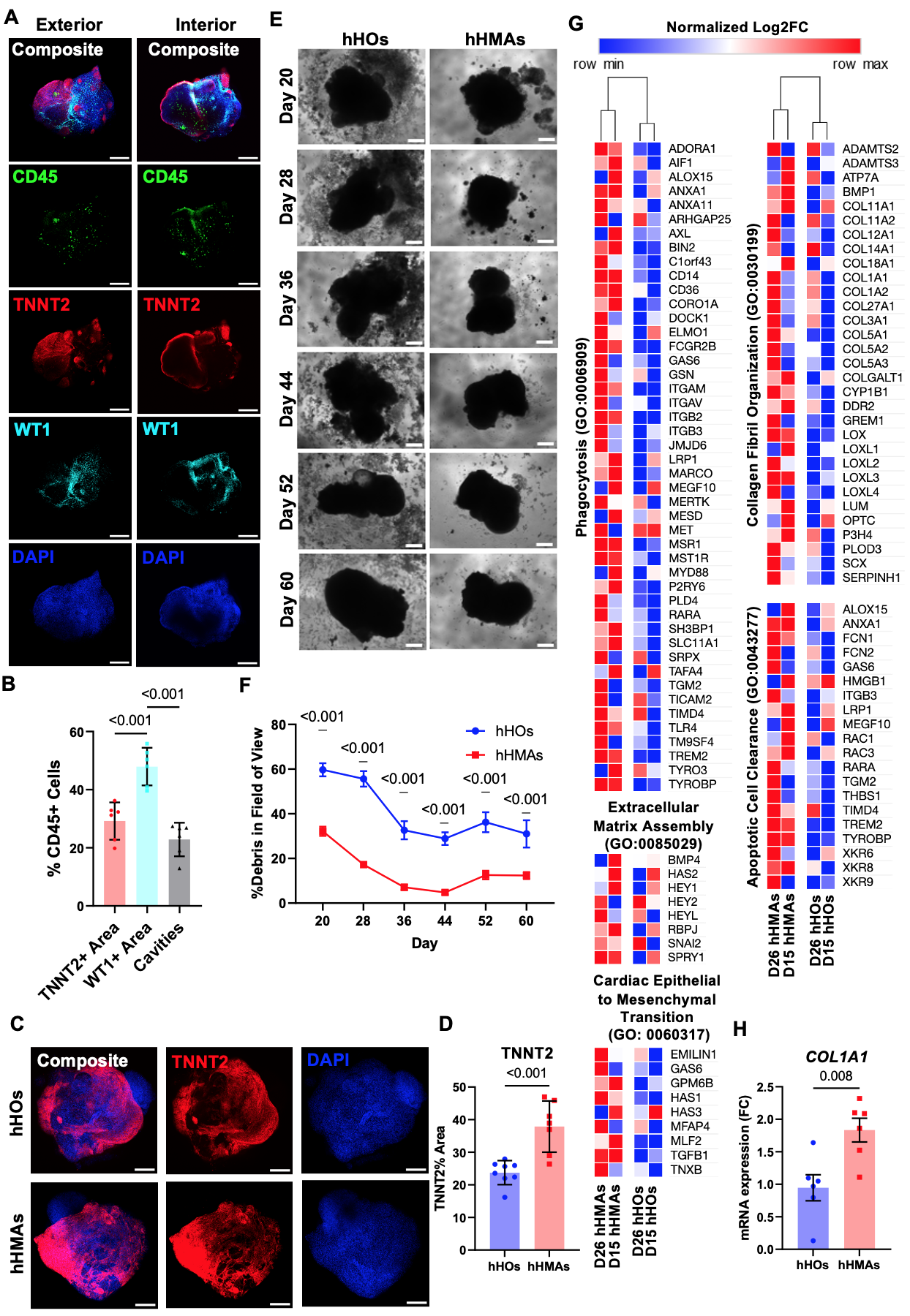

**Supplementary Figure 5. Autologous cardiac tissue-resident MPs associate more so with WT1+ cells than TNNT2+ cells, promote efferocytosis, and extracellular matrix assembly.**

(A) Representative confocal IF image for DAPI (blue), CD45 (green), WT1 (cyan) and TNNT2 (red), displaying the exterior (left) and interior (right) of day 20 hHMAs at low magnification. n = 6. Scale Bar = 200µm.

(B) Quantification of the proportion of total CD45+ cells found in TNNT2+ regions, WT1+ regions, and cavities of day 20 hHMAs averaged across 5 z-slices per hHMA. n ≥ 5. For graph: Value = mean ± s.e.m., 1-way ANOVA multiple comparison test.

(C) Representative low-magnification confocal IF image for DAPI (blue) and TNNT2 (red) in day 26 hHOs (top row) and hHMAs (bottom row). n ≥ 7. Scale Bar = 200µm.

(D) Quantification of TNNT2+ area as a percentage of total area in confocal IF images of hHOs and hHMAs averaged across 5 z-slices per organoid. n = 6. Value = mean ± s.e.m., Student’s t-test.

(E) Representative phase-contrast microscopy images depicting individual hHOs and hHMAs at day 20, 28, 36, 44, 52, and 60. n = 8. Scale = 200µm.

(F) Quantification of the background positive signal as a percentage of total area (Debris Area + Organoid Area) in the field of view of phase-contrast microscopy images in B, which we defined as Debris. Value = mean ± s.e.m., 2-way ANOVA multiple comparison test.

(G) Heatmap depicting normalized log2 fold change for genes related to phagocytosis, apoptotic cell clearance, collagen fibril organization, extracellular matrix assembly, and cardiac epithelial to mesenchymal transition in day 15 hHOs, day 26 hHOs, day 15 hHMAs, and day 26 hHMAs. Red correlates to maximum relative expression and blue correlates to minimum relative expression.

(H) RT-qPCR analyses on bulk RNA from day 20 hHMAs for COL1A1. n ≥ 6. Value = mean ± s.e.m., Student’s t-test.

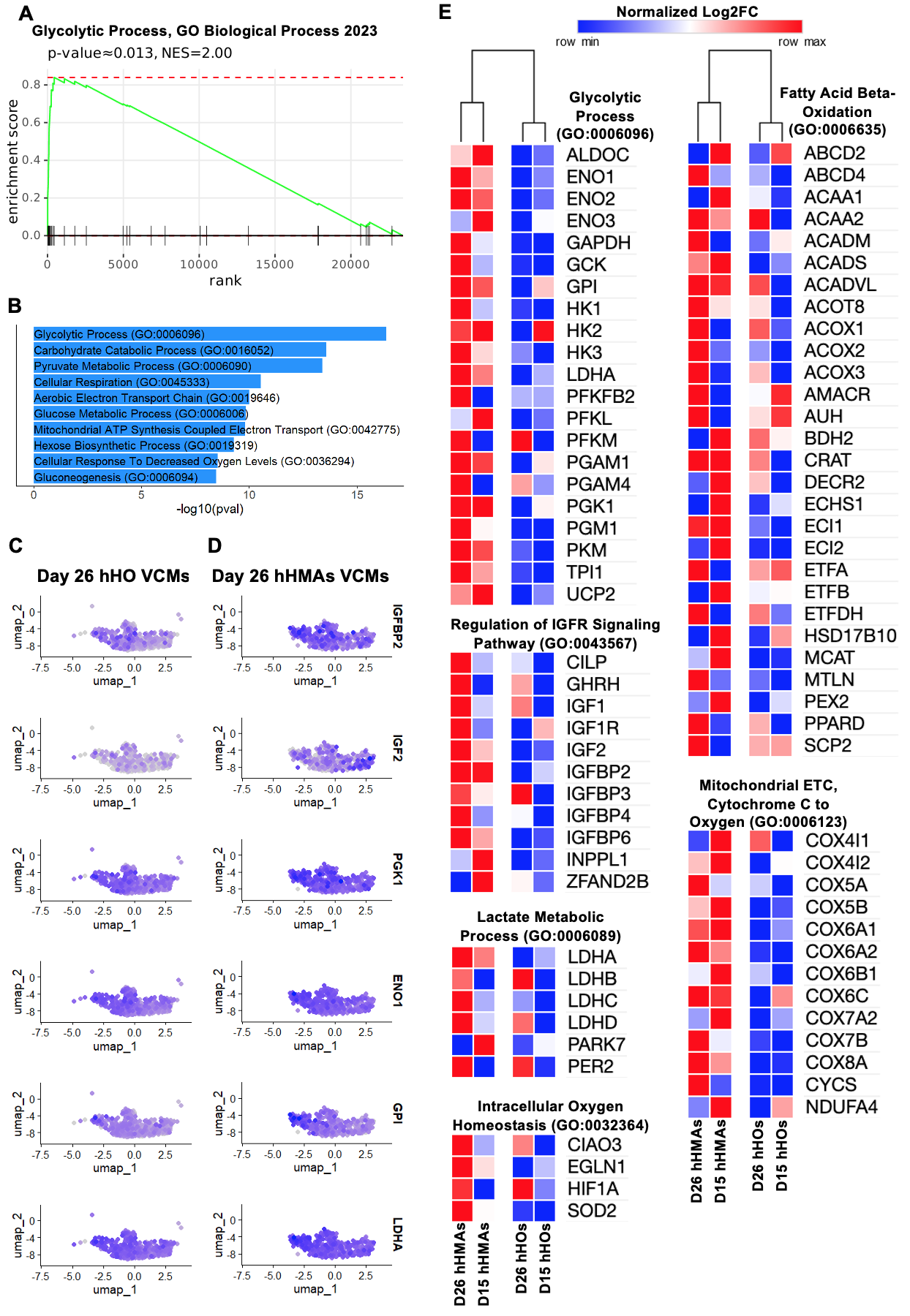

**Supplementary Figure 6. MPs enhance catabolic processes in hHMAs.**

(A) Gene set enrichment plots of differential gene expression between hHMAs and hHOs for Glycolytic Process, GO Biological Process 2023.
(B) Top 10 Gene ontology processes of 250 differentially expressed genes in MPs in Day 26 hHMAs versus Day 26 hHOs.

(C – D) Feature plots displaying upregulated genes related to glycolysis in VCMs of day 26 hHOs (C) and day 26 hHMAs (D). Color intensity represents the relative value of gene expression per gene.

(E) Heatmap depicting normalized log2 fold change for genes involved in glycolysis, IGFR signaling pathway, mitochondrial electron transport cytochrome C to oxygen, fatty acid beta-oxidation, lactate metabolic processes, and intracellular oxygen homeostasis in day 15 hHOs, day 26 hHOs, day 15 hHMAs, and day 26 hHMAs. Red correlates to maximum relative expression and blue correlates to minimum relative expression.

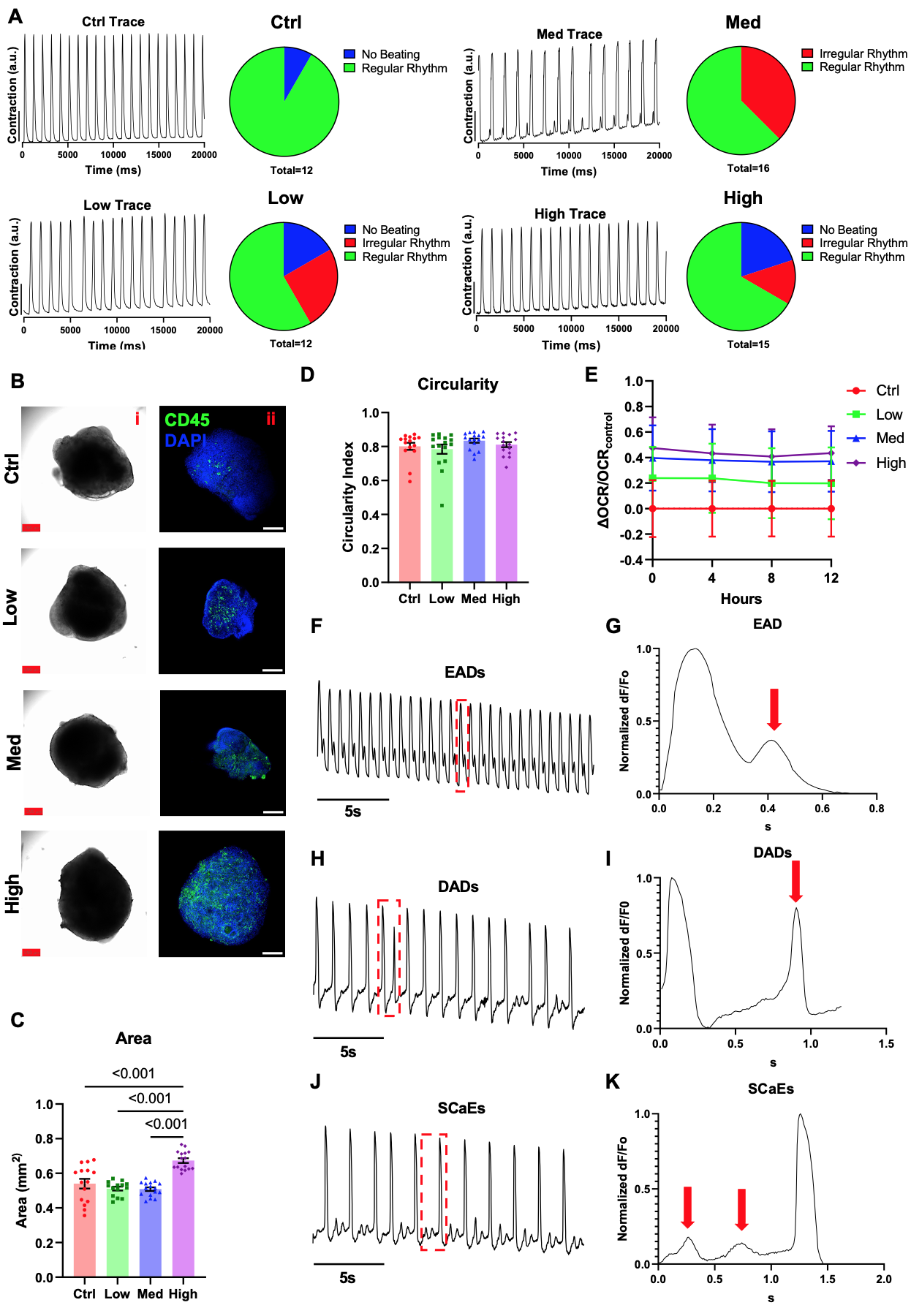

**Supplementary Figure 7: Effects of pro-inflammatory cytokines on hHMA action potentials, contractility, hHMA morphology, and oxygen consumption rate.**

(A) Representative muscle motion traces from phase-contrast microscopy videos of day 64 Ctrl, Low, Med, and High hHMAs. Pie charts categorize the hHMAs as either beating, irregular, or regular. n ≥ 12.
(B) Representative phase-contrast images (i) and immunofluorescence confocal microscopy images (ii) staining for CD45 (green) and DAPI (blue) of Ctrl, Low, Med, and High day 64 hHMAs. Scale bar = 200µm. n ≥ 15.
(C) Quantification of area (mm^2^) of day 64 Ctrl, Low, Med, and High hHMAs. n ≥ 15. Value = mean± s.e.m., 1-way ANOVA multiple-comparison test.
(D) Quantification of circularity (circularity index) of day 64 Ctrl, Low, Med, and High hHMAs. n ≥ 15. Value = mean± s.e.m., 1-way ANOVA multiple-comparison test.
(E) Normalized oxygen consumption rate of day 64 hHMAs from each condition measured over 20 hours. n ≥ 6. Value = mean± s.e.m., 1-way ANOVA multiple-comparison test.
(F) Representative FluoVolt trace depicting EADs in hHMAs. Dashed-red box displays an EAD seen in G.
(G) Representative FluoVolt trace depicting an EAD from the dashed-red box in F.
(H) Representative FluoVolt trace depicting DADs in hHMAs. Dashed-red box displays a DAD seen in I.
(I) Representative FluoVolt trace depicting a DAD from the dashed-red box in H.
(J) Representative FluoVolt trace depicting SCaEs in hHMAs. Dashed-red box displays SCaEs seen in K.
(K) Representative FluoVolt trace depicting SCaEs from the dashed-red box in J.
