## Supplementary material for "Human heart assembloids with autologous tissue-resident macrophages recreate physiological immuno-cardiac interactions": Key resources table

| **Antibody** | **Application** | **Provider** | **Catalog Number** | **Link** |
| --- | --- | --- | --- | --- |
| **Anti-Human TNNT2, Mouse** | IF | abcam | ab8295 | <https://www.abcam.com/products/primary-antibodies/cardiac-troponin-t-antibody-1c11-ab8295.html> |
| **Anti-Human TNNT2, Rabbit** | IF | abcam | ab45932 | <https://www.abcam.com/products/primary-antibodies/cardiac-troponin-t-antibody-ab45932.html> |
| **Anti-Human VCAM1, Mouse** | IF | DHSB | P3C4 | <https://sb-dshb.biology.uiowa.edu/P3C4> |
| **Anti-Human CD45, Rat** | IF | Invitrogen | MA5-17687 | <https://www.thermofisher.com/antibody/product/CD45-Antibody-clone-YAML501-4-Monoclonal/MA5-17687> |
| **Anti-Human NFATC1, Mouse** | IF | DHSB | 7A6 | <https://dshb.biology.uiowa.edu/7A6> |
| **Anti-Human CD14, Rabbit** | IF | SinoBiological | 205439-T02 | <https://www.sinobiological.com/antibodies/cd14-205439-t02> |
| **Anti-Human CD68, Mouse** | IF | BIORAD | MCA1815T | <https://www.bio-rad-antibodies.com/monoclonal/human-cd68-antibody-514h12-mca1815.html?f=s%2Fn> |
| **Anti-Human CD163, Mouse** | IF | BIORAD | MCA1853 | <https://www.bio-rad-antibodies.com/monoclonal/human-cd163-antibody-edhu-1-mca1853.html?f=purified> |
| **Anti-Human Connexin-43, Rabbit** | IF | Millipore Sigma | C6219-25UL | <https://www.sigmaaldrich.com/US/en/product/sigma/c6219> |
| **Anti-Human WT1, Rabbit** | IF | abcam | Ab89901 | <https://www.abcam.com/en-us/products/primary-antibodies/wilms-tumor-protein-antibody-can-r9ihc-56-2-ab89901> |
| **Anti-Human MYL3, Mouse** | IF | Santa Cruz Biotechnology | sc-47719 | <https://www.scbt.com/p/myl3-antibody-mlm508> |
| **Anti-Human NR2F2, Rabbit** | IF | abcam | ab211777 | <https://www.abcam.com/en-us/products/primary-antibodies/nr2f2-antibody-epr18443-ab211777> |
| **Anti-Human Calreticulin, Mouse** | IF | abcam | ab22683 | <https://www.abcam.com/en-us/products/primary-antibodies/calreticulin-antibody-fmc-75-ab22683> |
| **Anti-Human MERTK, Rabbit** | IF | abcam | ab52968 | <https://www.abcam.com/en-us/products/primary-antibodies/mertk-antibody-y323-ab52968> |
| **Anti-Human COL1A1, Mouse** | IF | DHSB | M-38 | <https://dshb.biology.uiowa.edu/M-38> |
| **Donkey anti-Rat IgG, AF488^TM^** | IF | Invitrogen | A-21208 | <https://www.thermofisher.com/antibody/product/Donkey-anti-Rat-IgG-H-L-Highly-Cross-Adsorbed-Secondary-Antibody-Polyclonal/A-21208> |
| **Donkey anti-Mouse IgG, AF647^TM^** | IF | Invitrogen | A-31571 | <https://www.thermofisher.com/antibody/product/Donkey-anti-Mouse-IgG-H-L-Highly-Cross-Adsorbed-Secondary-Antibody-Polyclonal/A-31571> |
| **Donkey anti-Mouse IgG, AF488^TM^** | IF | Invitrogen | A-21202 | <https://www.thermofisher.com/antibody/product/Donkey-anti-Mouse-IgG-H-L-Highly-Cross-Adsorbed-Secondary-Antibody-Polyclonal/A-21202> |
| **Donkey anti-Rabbit IgG, AF488^TM^** | IF | Invitrogen | A-21206 | <https://www.thermofisher.com/antibody/product/Donkey-anti-Rabbit-IgG-H-L-Highly-Cross-Adsorbed-Secondary-Antibody-Polyclonal/A-21206> |
| **Donkey anti-Mouse IgG, AF594^TM^** | IF | Invitrogen | A-21203 | <https://www.thermofisher.com/antibody/product/Donkey-anti-Mouse-IgG-H-L-Highly-Cross-Adsorbed-Secondary-Antibody-Polyclonal/A-21203> |
| **Donkey anti-Rabbit IgG, AF488^TM^** | IF | Invitrogen | A-21207 | <https://www.thermofisher.com/antibody/product/Donkey-anti-Rabbit-IgG-H-L-Highly-Cross-Adsorbed-Secondary-Antibody-Polyclonal/A-21207> |
| **FITC-Anti-Human CD45** | Flow | BD Pharmigen^TM^ | 555482 | <https://www.bdbiosciences.com/en-us/products/reagents/flow-cytometry-reagents/research-reagents/single-color-antibodies-ruo/fitc-mouse-anti-human-cd45.555482> |
| **AF647^TM^-Anti-Human CD14, Mouse** | Flow | BD Pharmigen^TM^ | 568242 | <https://www.bdbiosciences.com/en-us/products/reagents/flow-cytometry-reagents/research-reagents/single-color-antibodies-ruo/alexa-fluor-647-mouse-anti-human-cd14.568242> |
| **AF647^TM^-Anti-Human CD163, Mouse** | Flow | BD Pharmigen^TM^ | 568203 | <https://www.bdbiosciences.com/en-us/products/reagents/flow-cytometry-reagents/research-reagents/single-color-antibodies-ruo/Alexa-Fluor%252525E2%25252584%252525A2-647-Mouse-Anti-Human-CD163.568203> |
| **Anti-Human CD14, Microbeads** | MACS | Miltenyi Biotec | 130-050-201 | <https://www.miltenyibiotec.com/GB-en/products/cd14-microbeads-human.html#130-050-201> |

**Supplementary Table 1. List of Antibodies Used in Experiments.**

| **Gene** | **Application** | **Provider** | **Sequences (5’ 🡪 3’)** |
| --- | --- | --- | --- |
| ***HPRT1*** | RT-qPCR | IDT | **F: CGA GAT GTG ATG AAG GAG ATG G** |
|  |  |  | **R: TTG ATG TAA TCC AGC AGG TCA G** |
| ***PTPRC*** | RT-qPCR | IDT | **F: CGG CTG ACT TCC AGA TAT GAC** |
|  |  |  | **R: GCT TTG CCC TGT CAC AAA TAC** |
| ***CD14*** | RT-qPCR | IDT | **F: CTT GTG AGC TGG ACG ATG AA** |
|  |  |  | **R: TGC AGA CAC ACA CTG GAA G** |
| ***CD68*** | RT-qPCR | IDT | **F: ACG CAA CTG GCT CAA AGA** |
|  |  |  | **R: TCC CAA AGT GCT GGG ATT AC** |
| ***CSF1R*** | RT-qPCR | IDT | **F: ACC TCT TAG TCT CTG CCC TAT AC** |
|  |  |  | **R: CTG CCA CTT GGC TCA TTA CA** |
| ***AIF1*** | RT-qPCR | IDT | **F: ATG AGC CAA ACC AGG GAT TTA** |
|  |  |  | **R: CTG CTA TAT TTG GGA TCG TCT AGG** |
| ***CD74*** | RT-qPCR | IDT | **F: CAG ATG CAC AGG AGG AGA AG** |
|  |  |  | **R: CAG TTG CTC ATT GTT GGA GAT AAG** |
| ***MRC1*** | RT-qPCR | IDT | **F: GGACGTGGCTGTGGATAAAT** |
|  |  |  | **R:** **ACCCAGAAGACGCATGTAAAG** |
| ***AREG*** | RT-qPCR | IDT | **F: CCT TTA TGT CTG CTG TGA TCC T** |
|  |  |  | **R: CC TCA GCT TCT CCT TCA TAT TTC** |
| ***COL1A1*** | RT-qPCR | IDT | **F: CTA AAG GCG AAC CTG GTG AT** |
|  |  |  | **R: TCC AGG AGC ACC AAC ATT AC** |
| ***NLRP3*** | RT-qPCR | IDT | **F: GAA GAG GAG TGG ATG GGT TTA C** |
|  |  |  | **R: TCT GCT TCT CAC GTA CTT TCT G** |
| ***CASP1*** | RT-qPCR | IDT | **F: TCT ACC TCT TCC CAG GAC ATT A** |
|  |  |  | **R: GGA CTT TCA GTA CCC TTT CCT C** |
| ***IL-1B*** | RT-qPCR | IDT | **F: CAAAGGCGGCCAGGATATAA** |
|  |  |  | **R: CTGAATGTGGACTCAATCCCTAG** |
| ***LGALS3*** | RT-qPCR | IDT | **F: CCT CGC ATG CTG ATA ACA ATT C** |
|  |  |  | **R: CTC ATT GAA GCG TGG GTT AAA G** |
| ***RYR2*** | RT-qPCR | IDT | **F: GAA CGA GAG GTC AGC GAA TAA G** |
|  |  |  | **R: ACC GTC AGA ATG GAG AAG AAA G** |

**Supplementary Table 2. List of Primers Used in Experiments.**

| **Reagent** | **Provider** | **Catalog Number** | **Link** |
| --- | --- | --- | --- |
| **CHIR-99021** | Fisher Scientific | 44-231-0 | <https://www.fishersci.com/shop/products/chir-99021-tocris-2/442310> |
| **Activin A** | Fisher Scientific | 338AC010 | <https://www.fishersci.com/shop/products/r-d-systems-human-mouse-rat-activin-a-recombinant-protein-6/338AC010?searchHijack=true&searchTerm=338AC010&searchType=RAPID&matchedCatNo=338AC010> |
| **BMP4** | Fisher Scientific | PHC9534 | <https://www.fishersci.com/shop/products/human-bmp-4-recombinant-protein-20/PHC9534?searchHijack=true&searchTerm=PHC9534+&searchType=RAPID&matchedCatNo=PHC9534>+ |
| **Wnt-C59** | Fisher Scientific | NC0710557 | <https://www.fishersci.com/shop/products/wnt-c59-6/NC0710557> |
| **SCF** | R&D | 255-SC-010 | <https://www.rndsystems.com/products/recombinant-human-scf-protein_255-sc> |
| **FLT-3** | R&D | 308-FK-025/CF | <https://www.rndsystems.com/products/recombinant-human-flt-3-ligand-flt3l-protein_308-fk> |
| **M-CSF** | R&D | 216-MC-025 | <https://www.rndsystems.com/products/recombinant-human-m-csf-protein_216-mc> |
| **IL-3** | R&D | 203-IL-010 | <https://www.rndsystems.com/products/recombinant-human-il-3-protein_203-il> |
| **IL-1β** | R&D | 201-LB-010 | <https://www.rndsystems.com/products/recombinant-human-il-1-beta-il-1f2-protein_201-lb> |
| **LPS** | Millipore Sigma | L4516-1MG | <https://www.sigmaaldrich.com/US/en/product/sigma/l4516> |
| **IFN-γ** | R&D | 285-IF-100/CF | <https://www.rndsystems.com/products/recombinant-human-ifn-gamma-protein_285-if> |
| **FGF2** | R&D | 233-FB-010 | <https://www.rndsystems.com/products/recombinant-human-fgf-basic-fgf2-bfgf-146-aa-protein_233-fb> |
| **VEGF** | PrepoTech | 100-20 | <https://www.fishersci.com/shop/products/human-vegf-165-recombinant-protein-peprotech-20/p-7246045> |
| **GM-CSF** | Miltenyi Biotec | 130-093-862 | <https://www.miltenyibiotec.com/US-en/products/human-gm-csf.html#quality-grade=research-grade:size=50-ug> |
| **TPO** | Miltenyi Biotec | 130-095-745 | <https://www.miltenyibiotec.com/US-en/products/human-tpo.html#Quality-grade=Research-grade:size=25-ug> |
| **Thiazovivin** | Fisher Scientific | 507531 | <https://www.fishersci.com/shop/products/thiazovivin-greater99percent-1/507531#?keyword=507531> |
| **Relesr** | Fisher Scientific | NC2331323 | <https://www.fishersci.com/shop/products/relesr-100ml/NC2331323?searchHijack=true&searchTerm=NC2331323&searchType=RAPID&matchedCatNo=NC2331323> |
| **Accutase** | Fisher Scientific | NC9464543 | <https://www.fishersci.com/shop/products/accutase-500ml-bottles/NC9464543#?keyword=NC9464543> |
| **FluoVolt** | Fisher Scientific | F10488 | <https://www.fishersci.com/shop/products/fluovolt-membrane-potential-kit/F10488?searchHijack=true&searchTerm=F10488&searchType=RAPID&matchedCatNo=F10488> |
| **Fluo-4** | Fisher Scientific | F14201 | <https://www.fishersci.com/shop/products/fluo-4-am-cell-permeant/F14201?searchHijack=true&searchTerm=F14201&searchType=RAPID&matchedCatNo=F14201> |
| **NucBlue** | Fisher Scientific | R37605 | <https://www.fishersci.com/shop/products/molecular-probes-nucblue-live-readyprobes-reagent/R37605?searchHijack=true&searchTerm=R37605&searchType=RAPID&matchedCatNo=R37605> |
| **DAPI** | Fisher Scientific | EN62248 | <https://www.fishersci.com/shop/products/dapi-hoechst-nucleic-acid-stains/EN62248#?keyword=62248> |
| **RPMI 1640** | Fisher Scientific | 11-875-093 | <https://www.fishersci.com/shop/products/gibco-rpmi-1640-medium-41/11875093#?keyword=11875093> |
| **RPMI 1640, no glucose** | Fisher Scientific | 11-879-020 | <https://www.fishersci.com/shop/products/gibco-rpmi-1640-medium-no-glucose/11879020?searchHijack=true&searchTerm=11879020&searchType=RAPID&matchedCatNo=11879020> |
| **Essential 8 Flex** | Fisher Scientific | A2858501 | <https://www.fishersci.com/shop/products/essential-8-flex-medium-kit/A2858501?searchHijack=true&searchTerm=A2858501&searchType=RAPID&matchedCatNo=A2858501> |
| **B27 Supplement** | Fisher Scientific | A3582801 | <https://www.fishersci.com/shop/products/gibco-b-27-plus-supplement-50x/A3582801#?keyword=%2017504-044> |
| **B27 Supplement Minus Insulin** | Fisher Scientific | A1895601 | <https://www.fishersci.com/shop/products/b-27-supplement-minus-insulin-2/A1895601#?keyword=A1895601> |
| **StemPro-34 SFM** | Fisher Scientific | 10639011 | <https://www.fishersci.com/shop/products/stempro-34-sfm-1x/10639011?searchHijack=true&searchTerm=10639011%3B&searchType=RAPID&matchedCatNo=10639011%3B> |
| **Cardiomyocyte Dissociation Kit** | STEMCELL TECHNOLOGIES | 05025 | <https://www.stemcell.com/products/stemdiff-cardiomyocyte-dissociation-kit.html> |
| **L-Carnitine** | Millipore Sigma | 8400920025 | <https://www.sigmaaldrich.com/US/en/product/mm/840092> |
| **3,3’,5-Triiodo-L-thyronine** | Millipore Sigma | T2877-100MG | <https://www.sigmaaldrich.com/US/en/product/sigma/t2877> |
| **Oleic Acid** | Millipore Sigma | O3008-5ML | <https://www.sigmaaldrich.com/US/en/product/sigma/o3008> |
| **Linoleic Acid** | Millipore Sigma | L9530-5ML | <https://www.sigmaaldrich.com/US/en/product/sigma/l9530> |
| **Palmitic Acid** | Millipore Sigma | P0500-10G | <https://www.sigmaaldrich.com/US/en/product/sigma/p0500> |
| **Ascorbic Acid** | Fisher Scientific | A25215G | <https://www.fishersci.com/shop/products/l-ascorbic-acid-2-phosphate-sesquimagnesium-salt-hydrate-tci-america-2/A25215G?searchHijack=true&searchTerm=l-ascorbic-acid-2-phosphate-sesquimagnesium-salt-hydrate-tci-america-2&searchType=Rapid&matchedCatNo=A25215G> |
| **Glutamax Supplement** | Fisher Scientific | 35050-061 | <https://www.fishersci.com/shop/products/gibco-glutamax-supplement-2/35050061?searchHijack=true&searchTerm=35050-061&searchType=RAPID&matchedCatNo=35050-061> |
| **IGF-1** | Repligen | 10-1010-5 | <https://www.repligen.com/products/cell-culture-supplements/long-r3-igf-i> |
| **Glucose Solution** | Fisher Scientific | A2494001 | <https://www.fishersci.com/shop/products/gibco-glucose-solution/A2494001?searchHijack=true&searchTerm=A2494001&searchType=RAPID&matchedCatNo=A2494001> |
| **PBS** | Fisher Scientific | 10010049 | <https://www.fishersci.com/shop/products/pbs-ph-7-4-9/10010049?searchHijack=true&searchTerm=10010049&searchType=RAPID&matchedCatNo=10010049> |
| **Triton X-100** | Millipore Sigma | T8787-50ML | <https://www.sigmaaldrich.com/US/en/product/sigma/t8787> |
| **Paraformaldehyde** | VWR | 0215014601 CP-RF | <https://us.vwr.com/store/product/14512279/paraformaldehyde-powder> |
| **Bovine Serum Albumin** | Fisher Scientific | 50253966 | <https://www.fishersci.com/shop/products/bovine-serum-albumin-heat-s/50253966#?keyword=50253966> |
| **EDTA** | Millipore Sigma | E9884-100G | <https://www.sigmaaldrich.com/US/en/product/sial/e9884> |
| **Normal Donkey Serum** | Millipore Sigma | S30-100ML | <https://www.sigmaaldrich.com/US/en/product/mm/s30m> |

**Supplementary Table 3. Reagent List.**
