## Supplementary video descriptions for "Human heart assembloids with autologous tissue-resident macrophages recreate physiological immuno-cardiac interactions"

**Supplementary Video 1:** 3D reconstruction of OCT image of a representative hHO. Interconnecting sections were segmented in purple.

**Supplementary Video 2:** 3D reconstruction of OCT image of a representative hMO. Interconnecting sections were segmented in purple.

**Supplementary Video 3:** Video of Figure 6E. Scale Bar = 10µm.

**Supplementary Video 4:** Representative FluoVolt live-cell imaging video of day 64 Ctrl hHMOs in Figure 7B. Yellow color correlates to maximum signal and black correlates to a lack of signal. n ≥ 20. Scale Bar = 10µm.

**Supplementary Video 5:** Representative FluoVolt live-cell imaging video of day 64 Low hHMOs in Figure 7B. Yellow color correlates to maximum signal and black correlates to a lack of signal. n ≥ 20. Scale Bar = 10µm.

**Supplementary Video 6:** Representative FluoVolt live-cell imaging video of day 64 Med hHMOs in Figure 7B. Yellow color correlates to maximum signal and black correlates to a lack of signal. n ≥ 20. Scale Bar = 10µm.

**Supplementary Video 7:** Representative FluoVolt live-cell imaging video of day 64 High hHMOs in Figure 7B. Yellow color correlates to maximum signal and black correlates to a lack of signal. n ≥ 20. Scale Bar = 10µm.
